# Operationalizing climate risk in a global warming hotspot

**DOI:** 10.1101/2022.07.19.500650

**Authors:** Daniel G. Boyce, Derek P. Tittensor, Susanna Fuller, Stephanie Henson, Kristen Kaschner, Gabriel Reygondeau, Kathryn E. Schleit, Vincent Saba, Nancy Shackell, Ryan Stanley, Boris Worm

**Affiliations:** Department of Biology, Dalhousie University, Halifax, Canada; Bedford Institute of Oceanography, Fisheries and Oceans Canada, Dartmouth, Canada; United Nations Environment Programme World Conservation Monitoring Centre, Cambridge, UK; Oceans North, Halifax, Canada, B3J 1E6.; National Oceanography Centre, Southampton, UK; Department of Biometry and Environmental System Analysis, University of Freiburg, Freiburg, Germany; Institute for the Oceans and Fisheries, Changing Ocean Research Unit, University of British Columbia, Vancouver, British Columbia, Canada; NOAA Northeast Fisheries Science Center, Geophysical Fluid Dynamics Laboratory, Princeton University Forrestal Campus, Princeton, NJ 08540

**Keywords:** climate change, risk, vulnerability, species, biodiversity, fisheries, northwest Atlantic, management

## Abstract

There has been a proliferation of climate change vulnerability assessments of species, yet possibly due to their limited reproducibility, scalability, and interpretability, their operational use in applied decision-making remains paradoxically low. We use a newly developed Climate Risk Index for Biodiversity to evaluate the climate vulnerability and risk for ∼2,000 species across three ecosystems and 90 fish stocks in the northwest Atlantic Ocean, a documented global warming hotspot. We found that harvested and commercially valuable species were at significantly greater risk of exposure to hazardous climate conditions than non-harvested species, and emissions mitigation disproportionately reduced their projected exposure risk and cumulative climate risk. Of the 90 fish stocks we evaluated, 41% were at high climate risk, but this proportion dropped to 25% under emissions mitigation. Our structured framework demonstrates how climate risk can be operationalized to support short- and long-term fisheries objectives to enhance marine fisheries’ climate readiness and resilience.

## Introduction

Climate change is a major driver of change in marine ecosystems (1), with critical consequences for fisheries productivity (2) and human well-being (3). It is widely accepted that the successful management and conservation of living resources under climate change will require a comprehensive understanding of the differential vulnerability of species and ecosystems to global warming.

Climate Change Vulnerability Assessments (CCVAs) have been promoted as a critical component of marine management under climate change, particularly in protected areas (4,5) and fisheries management (6-8). However, many management agencies have not yet integrated climate change impacts and associated vulnerability assessments, despite the benefits and importance thereof (9-12). This situation is troubling as climate change can erode the effectiveness of traditional management approaches (13-15). According to the Food and Agriculture Organization of the United Nations, developing an enhanced understanding of the differential climate vulnerability of fisheries could increase the integration of climate adaptation measures into management worldwide (7), yielding improved outcomes in a changing ocean.

Despite their potential to support climate adaptation efforts (4-8), existing CCVA frameworks have various limitations that could help to explain their low incorporation into management settings (*e.g.* 9,11,12). With few exceptions (16-18), CCVAs often yield a single vulnerability value across species ranges, even though geographic variation in vulnerability is widely established (17-25) and critical to developing climate considered conservation strategies (4,5); they often rely on semi-quantitative expert opinions rather than being quantitatively derived from empirical data (6,17,26), providing a barrier to consistently tracking changing vulnerability through time and limiting their reproducibility; they infrequently evaluate all three component dimensions of vulnerability: exposure, sensitivity, and adaptivity (27); are very rarely spatially or taxonomically exhaustive (22); and almost exclusively rank and compare species vulnerabilities in dimensionless units (6,16–18,22,26,27), which is problematic, as stakeholders and associated structured decision-making frameworks often require explicit risk assessments on absolute rather than relative scales.

Here, we overcome these limitations by using the newly-developed Climate Risk Index for Biodiversity (CRIB) framework (28) to estimate the climate vulnerability and risk for marine species across their native geographic distributions in the northwest Atlantic Ocean under two contrasting greenhouse gas emission scenarios (SSP5-8.5: high emissions and SSP1-2.6: high mitigation) to 2100. The CRIB produces empirically rooted estimates that represent climate risk to the *in situ* persistence of species and the biotic intactness of the ecosystems they support, and it captures the likelihood of adverse consequences (29) at individual locations within species’ native geographic distributions and entire ecosystems. This framework was previously applied globally to ∼25,000 species at a coarser (1°) spatial scale (28). Here, we demonstrate its flexibility and potential to operationalize climate risk knowledge for 2,045 species, three large ecosystems, and 90 fish stocks at the higher (0.25°) spatial scales relevant to applied decision-making and facilitate its uptake into management processes and jurisdictions. Our analysis was undertaken within Canada’s exclusive economic zone (EEZ) and was geographically bounded by the Northwest Atlantic Fisheries Organization (NAFO) divisions, hereafter the area of study (AOS; Fig 1). This area of the Northwest Atlantic is a global warming hotspot (15,30) and thus provides an ideal test case to explore the operationalization and management implications of the CRIB. To tangibly illustrate these concepts, we explore how climate risk can be used to support and inform the conservation of three example stocks, thus enhancing traditional management strategies toward climate readiness (31).

**Fig 1.**
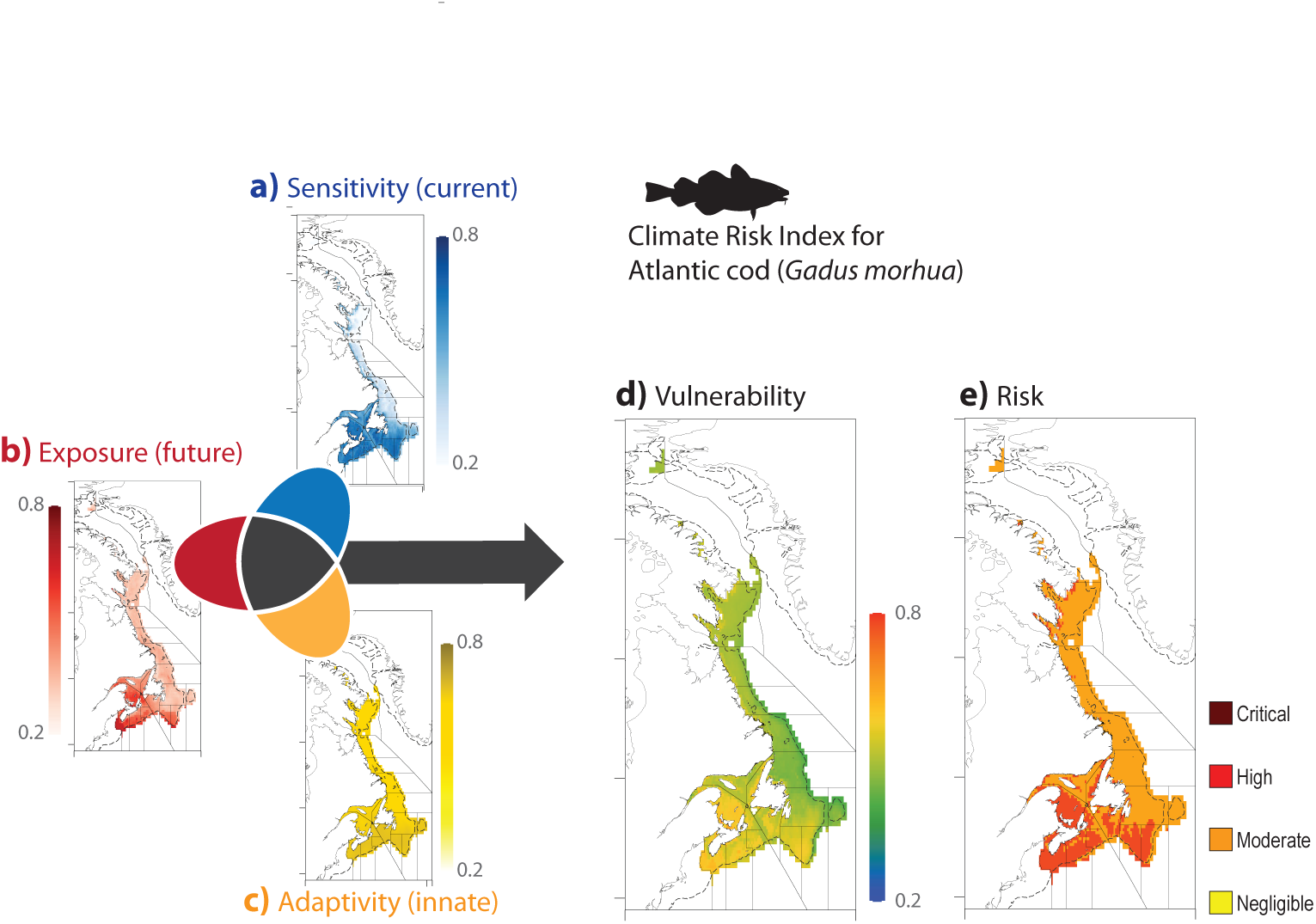
Spatially explicit assessment of climate vulnerability and risk for a single species, Atlantic cod (Gadus morhua). Within each grid cell (here 0.25°) across the native geographic distribution of cod within the AOS, 12 standardized climate indices are calculated and used to define the three dimensions of climate vulnerability: present-day sensitivity (a; blue), projected future exposure (b; red), and innate adaptivity (c; yellow). The dimensions are used to calculate cods’ climate vulnerability (d), and the relative vulnerability scores are translated into absolute climate risk categories for cod at all locations across its distribution (e).

## Results

### The climate risk index for biodiversity (CRIB)

For each species, publicly available and validated high-resolution data sources were used to calculate 12 climate indices in each 0.25° degree grid cell (∼24 km^2^ at 40° N) that comprise their native geographic distribution(28,32); (Methods section contains a description and rationale for the selection of indices). Fig 1 demonstrates how the CRIB framework calculates the indices for a single species, Atlantic cod (*Gadus morhua*), and integrates them to produce spatially explicit estimates of climate vulnerability and risk under a high mitigation scenario (SSP1-2.6). Cod experiences substantial variation in climate sensitivity (Fig 1a), exposure (Fig 1b) and adaptivity (Fig 1c) across its geographic distribution, with the climate exposure also varying considerably between emission scenarios. The cumulative climate vulnerability for cod (Fig 1d) is highest in nearshore and southerly locations (<50° N), where climate sensitivity and exposure tend to be higher and are only marginally offset by the higher adaptivity potential there. The relative climate vulnerability scores are then assessed against four ecologically rooted climate risk categories, enabling them to be interpreted on an absolute scale: negligible, moderate, high, and critical risk of hazardous climate impacts (Fig 1d). Thresholds that define the risk categories were independently identified in the 12 climate indices and were carried forward through the analysis. Climate risk thresholds were defined using a "thresholding" approach that is comparable to the definition of extinction risk applied by the IUCN Red List of species (33), the reasons for concern (RFC) framework adopted by the Intergovernmental Panel on Climate Change (IPCC)(29), and limit reference points that define the status of fisheries and are used to set harvest strategies (34). For each of the 12 climate indexes, we used ecological knowledge to set thresholds that define negligible, moderate, high and critical risk, thus enabling us to translate the dimensionless vulnerability scores to absolute risk categories (28). This translation of relative vulnerability into absolute risk is a critical step toward using climate vulnerability estimates in applied management settings such as fisheries, where precise, objective determinations of risk can facilitate tactical decision decision-making. Cod predominantly experiences moderate to high climate risk across its geographic distribution yet is at critical risk at some locations in the southern Gulf of St. Lawrence (GSL; Fig 1e). Within the CRIB definition of climate risk, cod will almost certainly experience negative climate change impacts at these locations under the high emissions scenario. However, emissions mitigation reduced the spatial extent of climate vulnerability and risk for cod; under high emissions, cod were at critical climate risk across 4% of their geographic distribution, but this dropped to 1% with high mitigation.

Almost all (98%) of the assessed species were animals, with Chordates (n=515) and Molluscs (n=506) each comprising ∼ one-quarter of the assessed species (25%), while Cnidarians and Arthropods made up 19% and 15% respectively (Fig S1a). More species were present in nearshore (<200m depth) and southern portions of the AOS relative to oceanic and northern locations (Fig S1b). The average body length of assessed species also varied spatially, with more large-bodied species being distributed in oceanic and some high latitude locations (Fig S1c).

### Climate risk for ecosystems and species

The vulnerability maps for all species were superimposed to evaluate geographic patterns of climate risk for ecosystems across the AOS under both emission scenarios to 2100. A greater fraction of species were at high or critical climate sensitivity risk in nearshore locations, with risk declining sharply outside the 100 m isobaths (Fig 2a). A greater fraction of species were climate-sensitive at lower latitudes (<50° N), particularly in the Gulf of St. Lawrence (GSL; Fig S2a). The GSL is a climate change hotspot (11,35), having experienced rapid surface warming (1900-2019), increased hypoxia (1984-2016) and acidification (35), declining sea ice extent (1979-2017), and high cumulative human impacts (36). The proportion of species at high climate adaptivity risk increased with latitude, peaking in the Arctic (>65° N). The geographic patterns in the proportion of species exposed to hazardous climate conditions under the low and high emission scenarios were similar (Fig S2c, S2d). However, more species were at risk of exposure under high emissions, with the proportion of highly exposed species increasing with latitude; some regions in both the Arctic Ocean and GSL had 100% of their species at risk of exposure. Overall, under both emission scenarios, the proportion of species at high or critical climate change risk tended to be higher in closer proximity to coastlines (stepwise increases at <500 m and <2000 m isobaths), particularly in the GSL and on the Grand Banks (Fig 2a). A larger fraction of species were at risk at high latitudes (>60° N), where the variability in climate risk scores was also higher. Under high emissions, most nearshore ecosystems had between 15% and 50% of their species at risk of adverse climate impacts, with some high latitude nearshore cells having >75% at risk. With emissions mitigation, the proportion of at-risk species declined at almost all locations, with mitigation benefits being most substantial at nearshore and high (>60° N) latitudes (Fig 2b).

**Fig 2.**
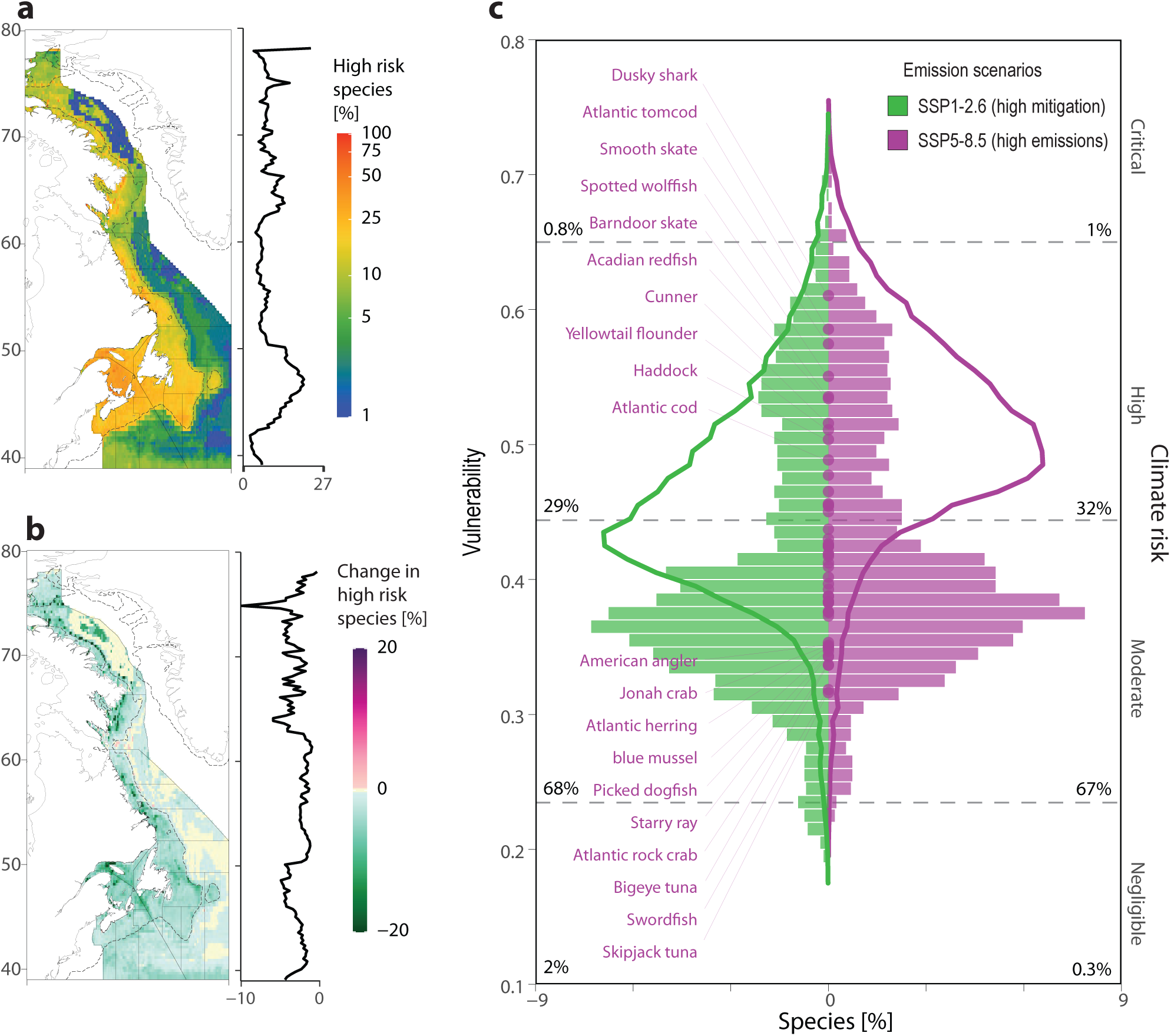
Climate vulnerability and risk for marine ecosystems and species. (a-The proportion of species at high or critical climate risk under high emissions in each grid cell to 2100. b) The difference in the proportion of species at high or critical climate risk in each grid cell under the low, relative to high, emissions scenario to 2100. Black lines denote the NAFO divisions, and the dotted line is the 200m isobath. (a-b) The variations along latitude are displayed in the right margins. c) Shading depicts the numerical densities of the vulnerability scores for all assessed species across the area of study under contrasting emission scenarios to the year 2100. Smoothed lines are the vulnerability scores for the global species pool (n=∼25,000 species) estimated in Boyce et al. (28). Gray dotted lines represent climate risk categories. Colours represent the emission scenario (low emissions=green; high emissions=purple).

On average, species were at high or critical climate risk across 33% and 29% (ranges:0-100%) of their native geographic distributions, under the low and high emission scenarios, respectively. Regardless of the emission scenario, when species’ vulnerability scores were averaged across their geographic distributions, over two thirds (67-68%) were at moderate risk of adverse climate change impacts, over one quarter (29-32%) were at high risk, and <2% were at negligible or critical risk (Fig 2c).

### Climate risk in a socioeconomic context

The status of marine species is closely tied to the health, culture, economy, and well-being of coastal communities in Atlantic Canada (11). To understand climate risk in a socioeconomic context, we evaluated the risk for (*i*) commercial species with an associated landed economic value and (*ii*) harvested species that are fished for subsistence, profit, or cultural reasons. Commercial species (n=17) collectively account for 87-91% of the total landed value in the region (2010-2019); of these American lobster (*Homarus americanus*), snow crab (*Chionoecetes opilio*), northern shrimp (*Pandalus borealis*), and sea scallop (*Placopecten magellanicus*) accounted for 84% (Fig 3a). The climate risk for these commercial species was first examined spatially by calculating the geographic distribution of aggregate (summed) species value (*LV_a_*; Fig 3b) and of climate risk and exposure in each grid cell. Intersecting those locations with the highest *LV_a_* and climate risk identifies priority ecosystems that collectively support many high-value species that are also at climate risk. Our analysis shows that *LV_a_* and aggregate species climate risks are positively associated under high (r=0.96) and low (r=0.94) emission scenarios. Climate risk and *LV_a_* predominantly intersect at lower latitudes (<50° N), in nearshore locations, and across known productivity hotspots such as the southern Grand Banks, Georges Bank, the Bay of Fundy, and offshore submarine banks (Fig 3c), where fishing tends to be concentrated. The intersection of projected climate exposure and *LV_a_* was similar but was to be more concentrated in the Gulf of St. Lawrence and less on the Grand Banks (Fig 3d)

**Fig 3.**
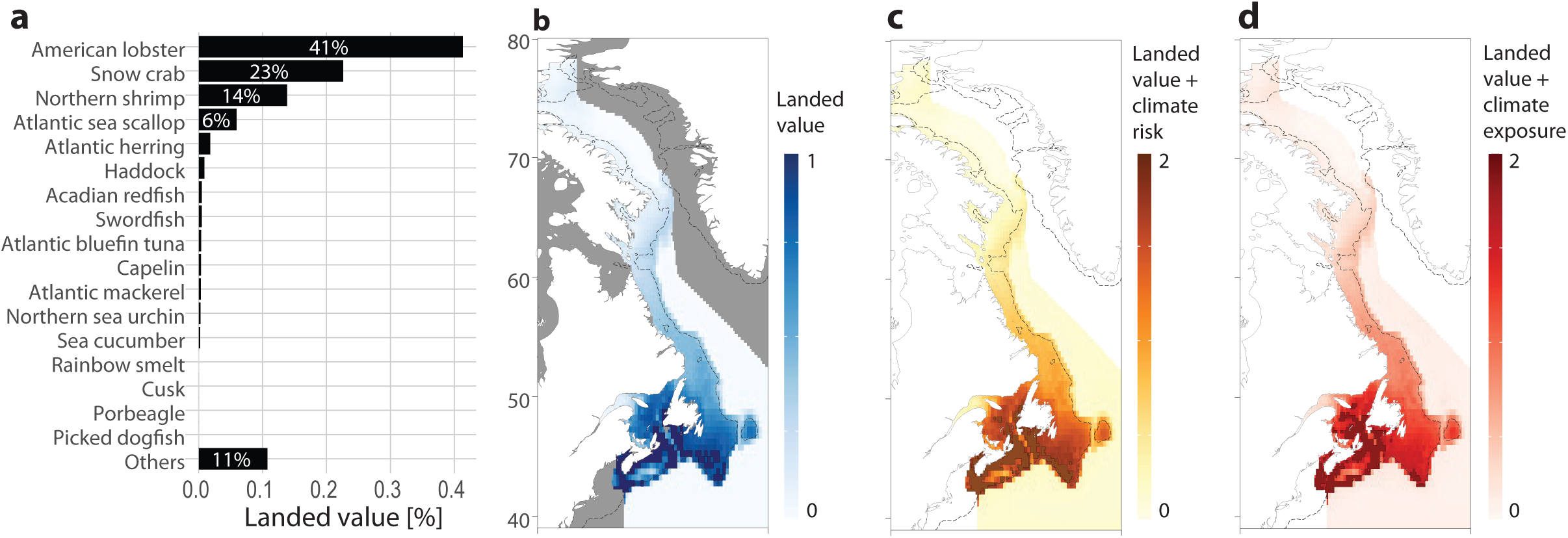
Climate risk for commercial species. (a) Contribution of the 17 commercial species to the total landed value of all seafood in Atlantic Canada (2010-2019). (b) Spatial patterns in aggregated landed value across the AOS; Darkest blue show areas where many species exist that collectively have a high landed value. (c-d) The intersection of aggregated landed value, (c) climate risk, and (d) exposure. (c-d). Darker shading show areas that have many species that collectively have high landed values and climate risk. (b-d) The darkest shading and outlined cells depict the top 5% of the values.

At the species level, the cumulative climate risk was comparable for commercial and non-commercial species, yet commercial species had a significantly greater risk of exposure to future climate impacts (Fig S3). The proportion of commercial species at risk of climate exposure was over three times higher (70%) than non-commercial species (22%). Differences in the spatial extent of climate exposure were also apparent. Non-commercial species were at high exposure risk across 12% of their distributions under low emissions and 23% under high emissions, so mitigation yielded an overall 11% reduction in the spatial extent of their climate exposure risk. However, commercial species were at high exposure risk across 22% of their distributions under low emissions and 75% under high emissions. Thus, the emissions pathway has a disproportionately strong impact on the climate exposure reduction of commercial species; mitigation reduced the spatial extent of their climate exposure risk by an average of 53%, over four times the reduction experienced by the non-commercial species. Further, some of the most considerable emission mitigation benefits - reductions in the proportion of their distributions at high exposure risk - were observed in the highest value species, including American lobster (-97%), sea scallop (-94%), northern shrimp (-86%), and snow crab (-68%); (Fig S3).

Harvested species include commercial species and those used for subsistence or cultural reasons (n=52). The climate risk of harvested species was not statistically different from those for non-harvested species (P>0.05; Fig 4c), yet harvested species tended to have higher climate sensitivity and exposure and lower adaptivity (P<0.05) than non-harvested species (Fig 4a, 4d). The proportion of harvested species at risk of exposure to projected hazardous climate conditions was roughly three times higher (62%) than non-harvested (22%). Under high emissions, harvested species were at high or critical risk of climate exposure across 63% of their native geographic distribution, compared to 23% for non-harvested species, on average; these values dropped to 21% and 11%, respectively, under the high mitigation scenario.

**Fig 4.**
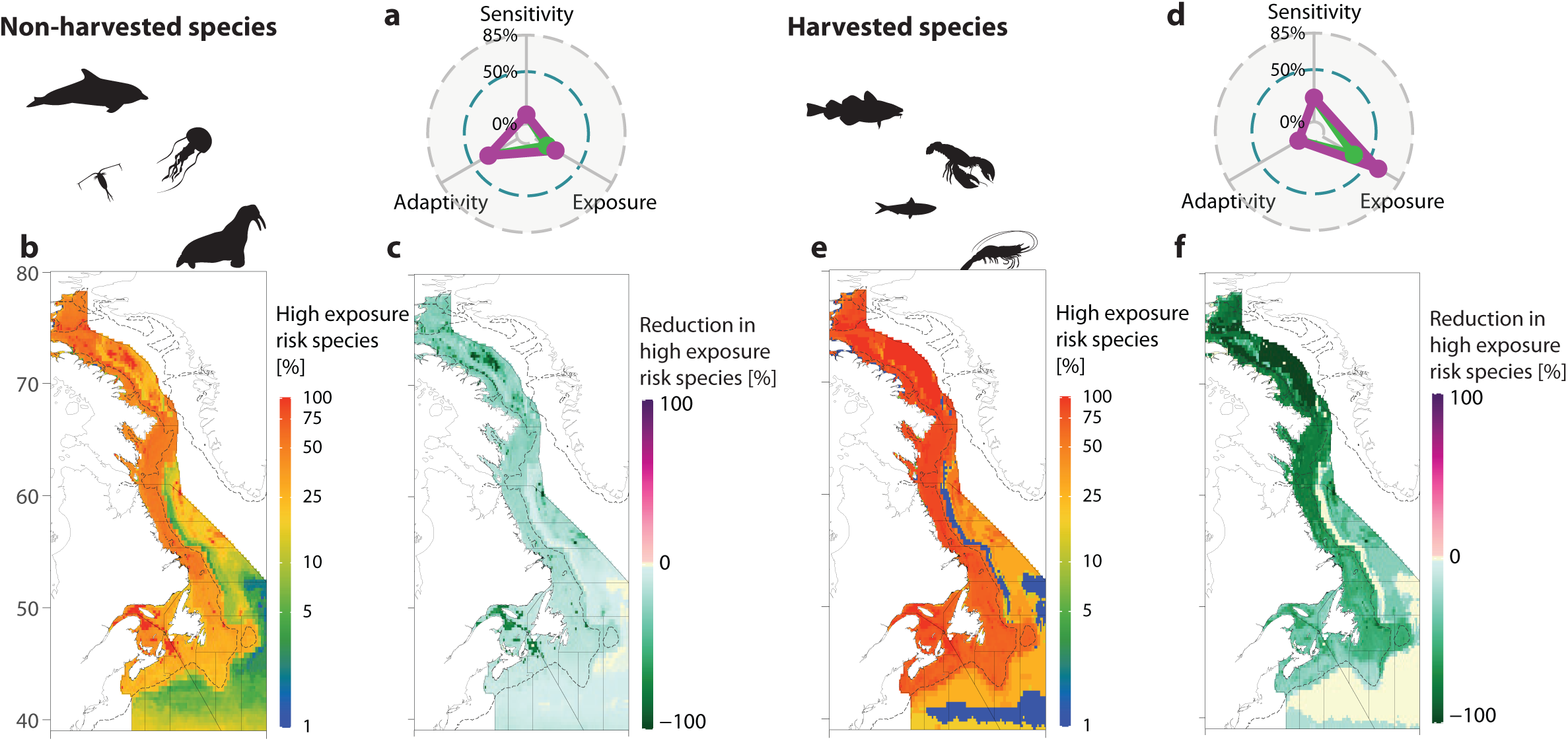
Comparative climate risk for harvested and non-harvested species. (a, d) Points depict the proportions of at-risk species (high or critical) according to their sensitivity, exposure, and adaptivity to climate change, under contrasting emission scenarios. b, e) The proportion of species at high or critical risk of exposure to hazardous climate conditions under high emissions in each grid cell to 2100. (c, f) The difference in the proportion of species at high or critical climate exposure risk in each grid cell under the low, relative to high, emissions scenario to 2100. (b, c, e, f) Black lines denote the NAFO divisions, and the dotted line is the 200m isobath.

The notably higher climate exposure of commercial and harvested species is driven by their anticipated responses to emissions-driven climate changes. Harvested species tended to inhabit more southerly areas of rapid projected climate velocity (Fig S7) and experience larger projected losses in their thermally suitable habitat to 2100. For example, with high emissions, the average loss of thermally suitable habitat for harvested species was 50%, compared to 19% for non-harvested. Harvested species were also disproportionately climate-sensitive (23%) relative to non-harvested (9%), possibly due to their lower conservation statuses and tendency to inhabit highly impacted environments. However, counterbalancing their high sensitivity and exposure, harvested species were also more adaptive to climate changes. The proportion of harvested species at risk in their climate adaptivity was much lower (8%) than for non-harvested (33%). Harvested species’ higher adaptivity may partly be driven by their broader and more contiguous geographic distributions relative to non-harvested species and their greater exposure to climate fluctuations that may pre-adapt them to climate changes(37). Notwithstanding the disparities in their climate sensitivity, exposure, and adaptivity, there were no significant differences in the overall climate vulnerability of harvested and non-harvested species under either emission scenario (P>0.05).

### Operationalizing climate risk for fisheries

Under contrasting emission scenarios, species climate vulnerability and risk maps were intersected with fisheries management zones to evaluate climate risk for 90 fish stocks across the AOS. Of these stocks, 75 (83%) are directed fisheries, 14 (16%) are caught as bycatch, and one (2J3KL Winter skate; *Leucoraja ocellata*) is under a fishing moratorium. Under contrasting emission scenarios, each stock’s climate vulnerability and risk were evaluated using their spatial management unit areas. For example, the risk to Atlantic cod (*Gadus morhua*) stocks was calculated by superimposing the geographic boundaries of the fishery management domain on the climate vulnerability and risk maps for cod (Fig S4). For each cod stock (n=8), climate vulnerability and risk, as well as its dimensions (n=3) and indices (n=12), can be visualized spatially to identify high-risk locations (*e.g.,* Figs S4a, S4b) or averaged across the spatial management area to yield a single value for each stock (*e.g.,* Fig S4c).

Under high emissions, 37 stocks (41%) were at high climate risk, with 53 (59%) at moderate risk (Fig 5). With high mitigation, the number of high-risk stocks dropped to 25 (28%), with 65 (72%) being at moderate risk. The elasmobranch stocks (n=7) had the greatest range of vulnerability and risk scores, with smooth skate (*Malacoraja senta*) in the southern GSL (4T) being the most vulnerable and thorny skate (*Amblyraja radiata*) on the Scotian Shelf (4VWXs) being among the least vulnerable, overall; Neither are targeted for capture but are retained as bycatch in other fisheries. With low emissions, stocks of herring (*Clupea harengus*), capelin (*Mallotus villosus*), and northern shrimp emerged as the least climate-vulnerable, although most were still at moderate risk. The lower climate risk of some demersal stocks such as monkfish (*Lophius americanus*) and lumpfish (*Cyclopterus lumpus)* may be partly driven by their deeper distributions, which under the CRIB framework assigns them a lower climate sensitivity. While emission mitigation benefited all stocks, some (*e.g.,* northern shrimps and lobsters) benefitted greatly, while others (*e.g.,* 4VWX5 silver hake; *Merluccius bilinearis*) only minimally. In general, emission mitigation led to a greater reduction in the climate risk of harvested species than non-harvested (Figs 3b, 3c, 3e, 3f). Despite the importance of evaluating, ranking, and summarizing the climate risk of fish stocks, reporting a single risk value per stock can also obscure substantial variation in climate vulnerability that occurs across the spatial area of each stock (Fig S5). For example, under high emissions, the 4RST capelin stock in the GSL is at high climate risk overall, but the risk ranges from moderate to critical across the stock area. This spatial variation in risk can be highly relevant to species conservation in general but particularly for fisheries, as they are harvested and often managed in a spatially explicit manner. The notable spatial variation in climate risk within and across stocks emphasizes the importance and value of assessing climate risk taxonomically and spatially to identify both species and locations at risk.

**Fig 5.**
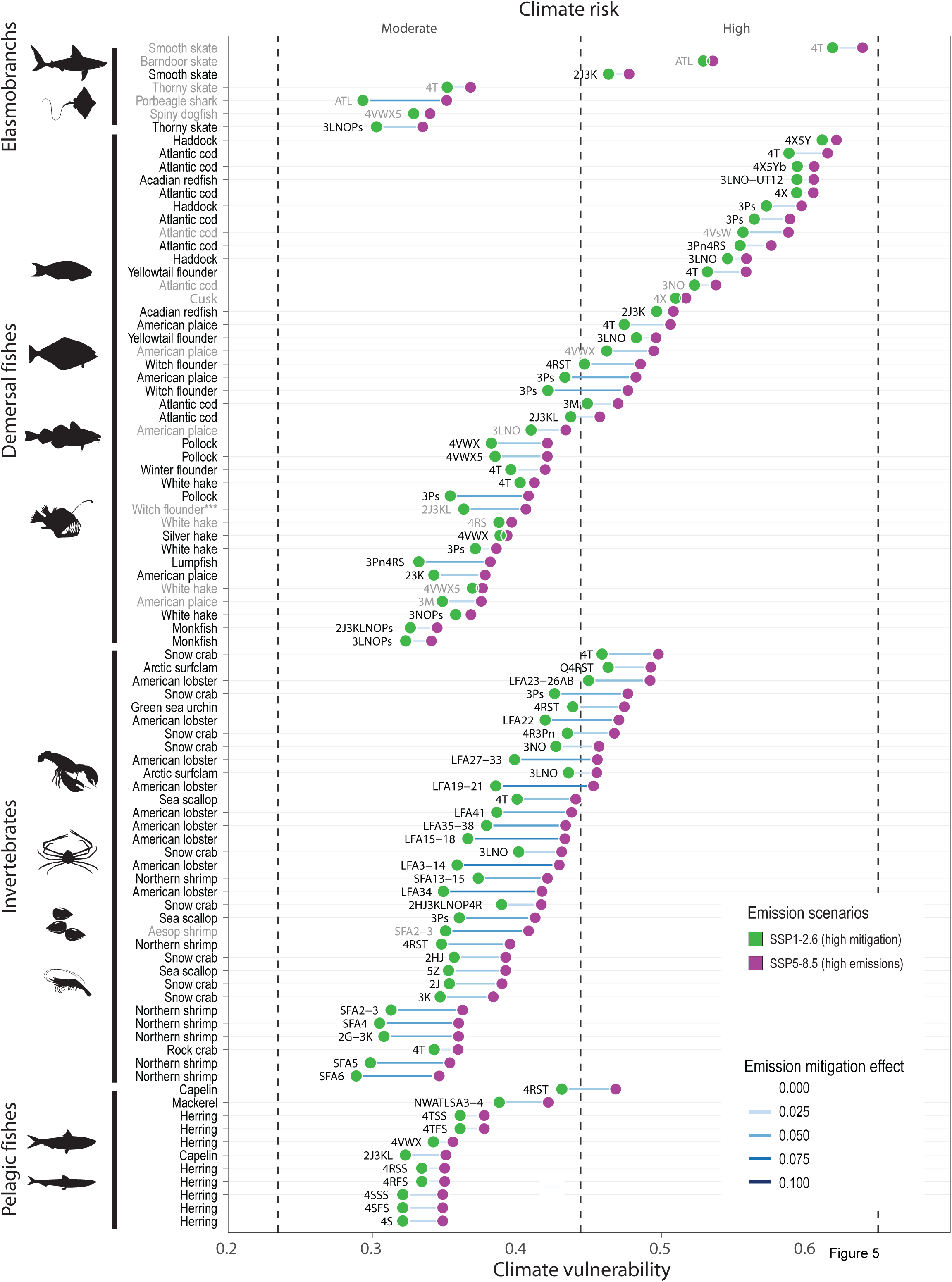
Climate risk and mitigation benefits for fisheries. Points are the average vulnerability scores for 95 stocks that operate across the AOS available within the RAM stock assessment database, estimated under contrasting emission scenarios to 2100. Coloured points represent the emission scenario (low emissions=green; high emissions=purple), and coloured lines the change in the average climate vulnerability of stocks with emission mitigation, where darker blue depicts larger emission mitigation effects. Black labels depict stocks for which there is a directed fishery and gray those fished as bycatch. ** indicates stocks under a fishing moratorium.

The assessed status of each fish stock within the Canadian Precautionary Approach (PA) framework (38) was evaluated against their climate risk to determine if stocks were systematically in double jeopardy, possessing both a high climate risk and poor management status. According to their reported stock biomass levels, the PA framework assigns each stock to one of four categories: critical, cautious, healthy, or uncertain. We calculated the climate risk for the 52 stocks assessed within the PA framework, targeted by fisheries operating within the AOS and had clearly defined and digitized stock management units. While the average climate vulnerability for fisheries that were reported in critical status was higher (0.48 SE ± 0.02; n=14) than for those in the cautious (0.42 SE ± 0.03; n=8), healthy (0.42 SE ± 0.02; n=15) or uncertain (0.41 SE ± 0.02; n=15), these differences were not statistically significant (p>0.05). However, stocks with critical status had a significantly greater climate exposure risk under high emissions than those assessed as healthy, cautious, or uncertain (p<0.05).

## Discussion

Our analysis suggests that irrespective of the emission scenario, and despite significant geographic and taxonomic variation, the climate risk for most marine species in Atlantic Canada is moderate or high, with fewer species having negligible or critical risk. However, on average, species in this global warming hotspot tended to be at lower climate risk when compared to the global species pool(28), where most of the ∼25,000 assessed species were at high climate risk (54-84%), with fewer species falling into the moderate risk category (13-44%; Fig 2c, colour lines). This difference suggests that species in the study area tend to have a lower overall climate risk than the global species pool and that their cumulative climate risk is less responsive to future emission pathways, on average. Several factors likely drive this difference. While current and projected surface warming rates in the northwest Atlantic are at the upper end of global warming trends (11,15,30), resident species tend to have broader thermal niches that render them generally less vulnerable to this rapid warming when compared to the global species pool. Furthermore, lower latitude ecosystems also experience rapid warming, and species there tend to possess narrower thermal niches on average (Fig S6); (39), which renders them generally more vulnerable. This result indicates that areas of high climate change or velocity may not necessarily be hotspots of ecological climate risk and emphasizes the critical importance of considering how species traits interact with the spatiotemporally dynamic environments they inhabit to define their climate risk robustly. Similar to the global analysis, climate risk across the northwest Atlantic was elevated in nearshore areas, likely because these locations tend to be more impacted by human activities (40), and species there tended to have poorer conservation statuses.

While harvested species have a similar overall climate risk to non-harvested species, the manifestation of risk differed significantly, with harvested species having a higher climate sensitivity and exposure and reduced adaptivity risk relative to non-harvested (Fig 3). The higher exposure of harvested and commercial species to projected hazardous climate conditions is especially notable. Under high emissions, harvested species were at high or critical risk of projected climate exposure across 63% of their native geographic distributions, compared to only 23% for non-harvested species, on average. At the species level and under high emissions, 62% of all harvested species are at high or critical risk of climate exposure, compared to 22% of non-harvested species. The higher climate exposure of harvested species is partly due to their greater projected loss of suitable habitat, which implies that they will experience disproportionate geographic displacement relative to non-harvested species across the AOS. Harvested species also tended to inhabit areas with higher cumulative impacts, which, along with their greater extinction risk and shallower depth distributions, renders them more sensitive to climate change than non-harvested species. The greater exposure and sensitivity of harvested and commercially valuable species could partly be explained by their greater frequency of occurrence in the southern portion of the AOS, and particularly in the GSL, relative to non-harvested species, particularly in the western portion (Fig S7), where climate changes and ecosystem stressors are elevated (11,40).

The higher climate exposure and anticipated geographic displacement of harvested species, particularly those of high commercial value (Fig S3), is concerning, given that there is presently low incorporation of climate change considerations and adaptation strategies in fisheries management across the AOS (9,11,12). For example, it was reported that climate change, as a general theme, arose in 11% of DFO research documents that form the scientific basis for fisheries management in Canada published between 2000 and 2020 and that the extent of climate change discussion in these documents was low (9,11). Similarly, it was found that incorporating climate change considerations into the management of marine conservation areas (*i.e.,* MPAs) lagged in Canada compared to other temperate jurisdictions (41). This apparent disconnect between climate risk and the management response raises the question of how our assessment of ecological climate risk can be integrated into the fisheries management process. While we can foresee several ways our climate risk analyses could be used to support species and ecosystem conservation initiatives (*e.g.* 28), our current focus is on fisheries sustainability.

Broadly, fisheries management represents the union of shorter-term (1-5 years) tactical objectives and actions and long-term (>5 years) strategic goals. Although sometimes viewed separately, these two spheres are inextricably linked: short-term tactical objectives centred on setting harvest rates to achieve present-day maximum sustainable yield also need to be set to help meet longer-term strategic goals such as stock recovery, sustainability, and in this case, robustness to climate variability and change. Climate risk knowledge could help support and inform management in both essential domains. Our climate risk analysis can aid in strategic planning by evaluating the effect of different emission pathways on economically valuable species and fisheries and how this might ultimately impact socioeconomically dependent communities. A Scenario Planning

Framework (42,43) could allow managers to explore outcomes of management strategies on species or areas of high climate risk. Understanding these tradeoffs between current socioeconomic development strategies and future ecological and economic consequences would allow responsible agencies to develop adaptation strategies and inform decisions about setting national strategies for climate mitigation as well as species-specific action plans. Climate risk can also help determine overarching strategic objectives and directions, such as developing a national climate change strategy for fisheries management (*e.g.* 44). For example, in the Canadian context, establishing an explicit climate change objective into Canada’s National marine conservation strategy (5), or explicitly incorporating climate change into legislation such as Canada’s Fisheries Act and/or Oceans Act. Additionally, using several approaches discussed here, climate risk tools could help support and inform fisheries management over shorter-term tactical scales. These examples embody the principle that healthy, sustainably fished populations exposed to fewer stressors are more resistant and resilient to climate change.

First, the climate risk analysis presented here can be incorporated fisheries and marine spatial planning by triaging fish stocks to identify and prioritize the species and locations most urgently in need of resources to enable climate-smart management and conservation (45). Rather than ranking fish stock vulnerability on relative scales that can be difficult to interpret, the CRIB produces explicit and spatially resolved risk assessments on absolute scales, thus facilitating risk communication to stakeholders and aiding decision-makers. Notably, the assessment framework used to define climate risk is conceptually similar to those in prolonged use by fisheries management agencies worldwide to identify stock status (*e.g.* 34). For example, in our analysis, the 4T smooth skate stock species of high conservation concern also had the highest climate vulnerability under both emission scenarios (Fig 5). This bycatch fishery is at high risk: it is critically exposed to projected climate changes, is critically sensitive to them, and has a moderate adaptivity (Fig 6a, 6b). The IUCN Red List (RL) classified this species as globally endangered (33), and the Committee on the Status of Endangered Wildlife in Canada (COSEWIC) has assessed the smooth skate populations in the Laurentian-Scotian region, encompassing the 4T stock, as special concern (46). In a climate triage system for fisheries, this stock would thus be a high priority for climate adaptation.

**Fig 6.**
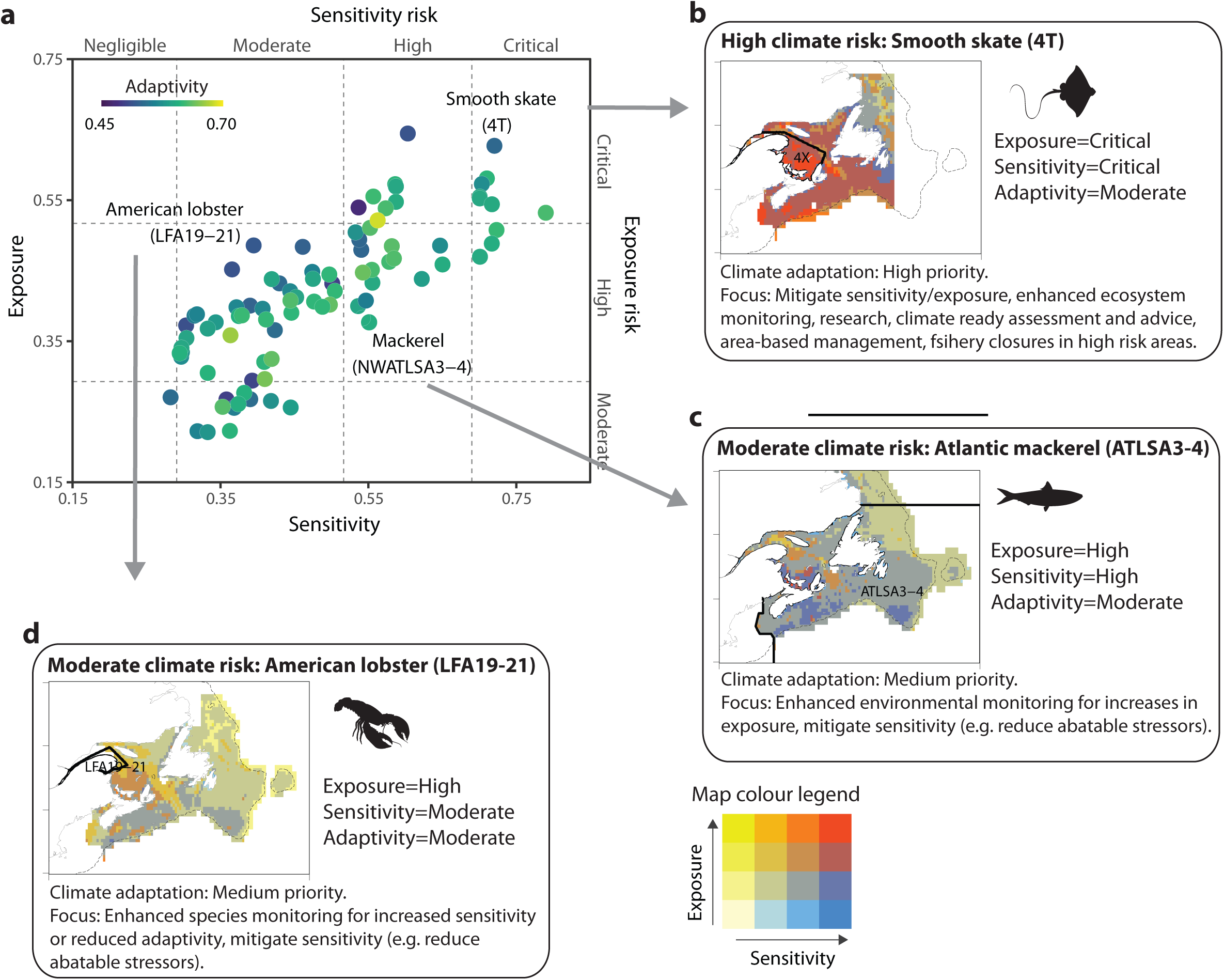
Operationalizing climate vulnerability and risk for fisheries. (a) Relationship between the scores and risk categories for climate sensitivity, exposure, and adaptivity of 95 stocks that operate across the AOS under the high emission scenario to 2100. Colours are the adaptivity scores (dark blue=low, yellow=high). (b-d) Examples of climate risk scenarios for fish stocks and how they can inform management priorities and decisions. Examples include b) Smooth skate (4T), a high-risk species, (c) Atlantic mackerel (ATLSA3-4), a moderate risk species but having higher sensitivity and lower exposure and adaptivity, d) Atlantic lobster (LFA19-21), a moderate risk species, but having higher exposure and lower sensitivity and adaptivity. (b-d) Climate sensitivity and exposure are displayed as colours: blues=high sensitivity-low exposure, yellows=high exposure-low sensitivity, reds=high sensitivity-high exposure. Stock areas are displayed as thick black lines and are labelled; the dotted line is the 200m isobath.

Resources that would support climate adaptation could include, for instance, pragmatic approaches that integrate climate change considerations into stock assessments, harvest advice, and decision-making (9,11,47,48). Approaches for doing so range from using climate risk as a modifier to the harvest advice to more detailed quantitative inclusion of climate variables (temperature, oxygen, pH) into the stock assessment process (49,50). Management strategy evaluations (MSE) can find candidate management strategies that are potentially robust to future climate scenarios, population and ecosystem dynamics and other uncertainties (51-53). Dynamic management can set harvest rates based on dynamic climate or ecological forecasts or respond in real-time to changing conditions (54). Adaptation resources could also include instituting flexible spatial protections where the stock is particularly at risk (4), targeted ecosystem monitoring for changes in climate vulnerability, and addressing factors affecting the sensitivity and adaptivity of smooth skate (*e.g.* reducing directed and ecosystem overfishing, bycatch, habitat destruction, pollution); (10). Since high natural mortality in the GSL, primarily by seals, is also thought to be impairing species recovery (46), understanding the influence of bycatch on their mortality and reducing any fisheries induced through bycatch reduction (*e.g.* targeted gear regulations and/or seasonal fishery closures, protected areas) could improve the climate adaptivity potential for 4T smooth skate, and species with similar climate risk. For transboundary species and stocks with high climate risk, priority should be placed on ensuring single species focused fisheries management is broadened to ecosystem-based fisheries management that can both predict and accommodate shifting geographical distributions across jurisdictional boundaries (55).

Second, while fisheries managers and stakeholders will ultimately determine the specific management strategies for each stock, climate change risk analyses (CCRAs) can help identify key overarching priorities, needed resources and focal actions under climate change. The dimensions of climate vulnerability and risk could be helpful in this context. For example, ATLSA3-4 stock for Atlantic mackerel (*Scomber scombrus*), a forage species of high ecological importance, is at moderate climate risk presently but possesses a higher climate sensitivity and lower exposure (Fig 6c). This species is listed as least concern by the Red List (33) yet has been assessed as falling into the critical zone of Canada’s Precautionary Approach Framework (56) and is scheduled to be assessed by COSEWIC for the first time in 2022. Further, there is serious concern over the conservation status of forage fishes such as mackerel that are of high ecological importance and support valuable upper trophic fisheries (57). While this stock is currently at moderate risk, increasing its exposure could increase its cumulative climate risk to high or critical. Consequently, this stock would be a medium priority for climate adaptation resources. Yet, actions could be taken to reduce climate sensitivity, such as developing interventions that minimize abatable stressors (*e.g.* directed and ecosystem overfishing, bycatch, pollution, habitat or ecosystem disruption) or through fisheries closures or climate-considered integrated marine spatial planning (4). For example, in 2022, commercial and bait fisheries for Atlantic Mackerel and Atlantic herring in the southern GSL were put under a moratorium to permit stocks to recover from historically low abundance. These measures can reduce stress and promote the recovery of these species and are one example of interventions that can be applied to reduce stressors. Enhanced environmental monitoring could be prioritized to detect further increases in the stock’s exposure to hazardous climate changes, for instance, by monitoring species’ thermal safety margins over time. As another example, the lobster stock in Lobster Fishing Areas (LFAs 19-21) is also at moderate climate risk overall (Fig 5). This species is not a conservation concern globally (33) and is not assessed by COSEWIC but has the highest economic value of any species in the region (8). This lobster stock possesses a higher climate exposure and lower sensitivity and adaptivity (Fig 6d), and an increase in its sensitivity or decline in adaptivity could shift its climate risk to high or critical. This stock would also be a medium priority for climate adaptation resources. However, given its economic importance, enhanced species monitoring could be conducted to identify increases in fisheries sensitivity (*e.g.,* lower conservation status, narrower thermal safety margins) or reductions in adaptivity (*e.g.,* smaller and more fragmented distribution) to climate change that could trigger climate-relevant management actions.

Third, the spatially resolved risk maps we present can help inform spatial fisheries management or area-based management tools by identifying areas where the species is particularly at risk or that function as climate refugia (58,59). This also applies to identifying spatial management units that will face extreme climate challenges. We report spatially explicit vulnerabilities that can help evaluate and manage the climate risk for transboundary and highly migratory fish stocks (55). They could also help pinpoint areas where an exploited species is in double-jeopardy, being at high climate risk at a location that also functions as critical essential habitat for its population (*e.g.,* spawning, summer feeding). Such areas could be priorities for spatial management measures such as seasonal fishery closures, spatial conservation measures (*e.g.,* marine protected areas or other effective area-based conservation measures) or enhanced monitoring.

Fourth, besides providing cumulative risk estimates for fisheries, our framework also contains more detailed information about the 12 individual aspects of their climate vulnerability. For example, the timing of climate emergence from species’ thermal niches (60,61) can provide a chronology of when a stock will first become exposed to hazardous climate conditions across its management area. For example, while the 4T smooth skate stock is projected to be exposed to a hazardous climate in 36 years (by 2056) on average (range=0-80 yrs.), the Atlantic mackerel stock is not expected to be exposed for another 74 years (by 2094) on average (range: 16-80 yrs.). Such information can aid in proactively developing timelines to institute adaptation resources in advance of those impacts and help understand the pace of climate impacts on fisheries.

Fifth, climate risk analyses can help reveal critical knowledge gaps and uncertainties that could erode management effectiveness under climate change. For instance, such knowledge gaps could arise for stocks identified as being at climate risk and thus in urgent need of climate adaptation resources (*e.g.,* climate-considered stock assessments) but for which these cannot be implemented due to resource constraints (*e.g.,* data, technical expertise, knowledge). In such situations, priority would be placed on filling in the missing knowledge or resources to facilitate climate adaptation.

Finally, our climate risk analysis can monitor changing vulnerability and risk of species, ecosystems, and fisheries over time in a standardized manner. The ability to track changing fisheries vulnerability in a spatially explicit, rapid, and cost-effective manner could provide crucial information about how fisheries are becoming more or less at risk and anticipate future climate risk outcomes.

Notwithstanding the potential utility of our climate risk analyses, the complexity of climate impacts requires any climate risk framework to make assumptions. We undertook this analysis using state-of-the-art earth system and species distribution models at the highest spatial resolution permitted by the input data. However, using higher-resolution data when it becomes available and incorporating additional climate variables (*e.g*., dissolved oxygen, pH, and sea ice) could improve the characterization of species distributions and climate risk, particularly in coastal locations. For instance, 10km^2^ may be an optimal resolution for resolving ocean climate processes across the northwest Atlantic Ocean (30), and evaluating risk at this scale could broaden its application, as climate-considered conservation and management strategies often require such fine-scale spatial information (4,5). Lastly, while our analysis evaluated thermal niches for each species, niches can also vary by population, and accounting for this source of variation in future could refine the climate risk for some species or stocks. Notwithstanding these caveats, our climate risk framework builds on existing approaches in several ways, including being spatially and taxonomically explicit, comprehensive, and risk-based. As shown here, the framework is designed to be flexible (*e.g.*, incorporate different data sources and new knowledge) and can potentially overcome several of its caveats as new information and knowledge become available.

## Conclusions

Our analysis agrees with a growing body of evidence suggesting that climate change affects fisheries’ productivity, resilience, and sustainability (2,62–64), with consequences for their sustainable management and conservation. Climate and marine ecosystem model projections suggest that this trend will continue into the foreseeable future (5,62,65,66), with socioeconomic repercussions (3,67). Such climate-driven changes underscore the critical need to increase the climate readiness of fisheries in Canada and elsewhere. This analysis demonstrates that the CRIB framework can further these objectives in several tangible and meaningful ways and supplement traditional fishery assessments. These include prioritizing climate adaptation resources, determining optimal management strategies, enhancing the effectiveness of spatial protections and marine spatial planning, identifying knowledge gaps and uncertainties, and tracking changing climate risk over time. While climate vulnerability and risk analyses are not a panacea to managing fisheries under climate change, our study demonstrates that if carefully undertaken, they can inform and complement existing assessment and management approaches and aid in determining how to prioritize and efficiently deploy limited climate adaptation resources for fisheries.

## Materials and methods

### Overview of CRIB design principles, methodology, and assessed species

The CRIB framework is fully described in Boyce *et al.* (28). Briefly, it incorporates information and features that are often required of climate risk assessments in applied settings, including i) it is spatially resolved, evaluating risk at all sites across species’ geographic distributions, ii) it produces relative vulnerability scores on a standardized scale and translates them into absolute risk categories, iii) it uses quantitative, well-validated, and publicly available data, thus ensuring reproducibility, iv) it is flexible and can be applied at differing spatial scales and biomes and can accommodate different information types, v) it is comprehensive, evaluating all three dimensions that define vulnerability and risk (68) using multiple assessment types (*e.g.,* trait-based, mechanistic, correlative); (26), vii) it assesses the statistical uncertainty (variability) of the vulnerability and risk scores, viii) it assesses the impacts of anticipated future climate conditions on species to facilitate decisions regarding emission mitigation, and ix) it is designed hierarchically, thus maximizing its flexibility and the information content (Fig 2).

The methodology used to calculate the CRIB has been described extensively in Boyce *et al*. (28). The 12 climate indices that define it capture climate change impacts that are generalized across species with varying life histories which are grounded in ecological theory, widely accepted and validated through peer review. The indices maximize parsimony and minimize redundancy and pseudoreplication: those that were easy to interpret and calculate were prioritized. The indices collectively include trait-based, correlative, and mechanistic information and incorporate abiotic, biotic, and human pressures acting across multiple biological organization levels from species to ecosystems. The indices integrate historical, present-day, and projected future information about species’ climate vulnerability and are calculated or obtained in their native units. The 12 climate indices are described in Table S1. The climate sensitivity indices include species’ thermal safety margins (16,23–25), vertical habitat variability and use (69-72), conservation status (73), and cumulative impacts (14,36,40,74–78). Climate exposure indices were based on ensemble climate projections and included the’ timing of climate emergence from species’ thermal niches (60,61,79,80), the extent of suitable thermal habitat loss (81-83), climate-related ecosystem disruption (60,84–87), and the projected climate velocity (88–90). Adaptivity indices included the species’ geographic range extent (69,88,90–94), geographic habitat fragmentation (17,95–99), maximum body length (17,26,97,100–104), and historical thermal habitat variability and use (17,105–108).

Each index was calculated from environmental or ecological data on a geographic grid across the native geographic distribution of the focal species, defined by the focal species’ traits and/or a mix of the two. This produces indices that are taxonomically (*e.g.,* each species) and geographically (*e.g.,* each grid cell) explicit. The indices are transformed to ensure they are on a standardized scale (0-1) across all species and locations. This step ensures that indices with different native units can be compared, normalized, and combined while at the same time ensuring that vulnerability can be calculated at different spatial resolutions or different points in time without losing information. Reference values and scaling functions were used to meet these criteria and are described in Boyce *et al.* (28). The 12 standardized climate indices are used to calculate three climate dimensions (sensitivity, exposure, and adaptivity), which ultimately define climate vulnerability and risk.

Species that do not live in the upper 100 m of the ocean are excluded from the analysis, and species with a maximum depth tolerance of more than 1000 m and a preference of more than 600 m are also excluded, as surface temperature may not well define the climate risk of these species. In order to verify this threshold, a sensitivity analysis was carried out in advance (28); (Fig S42 in ref. (28)). Seabirds were also excluded from the analysis because only a small part of their time is spent in surface water. However, mammals and endothermic fishes (*e.g.* tunas, billfishes) that can sometimes inhabit depths over 1000 m were not excluded; despite their ability to range into deeper waters, their distribution is often well explained by surface temperatures (109,110). Although it is impossible to assess their freshwater habitat, anadromous species (0.3% of the total) are included in the analysis as most species are primarily marine; freshwater habitats account for a minority of their total geographic distribution. Finally, guided by sensitivity analyses (Supplementary Fig 43 in ref. (28)), we restricted our analysis to species and cells containing all 12 indices and species that lacked at least one climate index in more than 10% of their native range were removed from the analysis.

### Data

Several data sources are identical to those used in a global implementation of CRIB (28) and are described therein. However, two alternative data sources were used to implement the analysis at the higher spatial resolutions within Canada. Firstly, instead of using climate projections at a global 1° resolution, they were instead obtained at a 25km^2^ native resolution and interpolated to 0.25° (25km^2^ at 40° N). Secondly, instead of using species’ global conservation status, where possible, we obtained species conservation status’ that were specifically relevant to different regions within Canada from the Wild Species General Status of Species in Canada reports(111). Further, species distributions and environmental data layers were acquired or re-gridded to a standardized 0.25° resolution. All data sources are described below and are listed in Table S2.

### Climate projections

An ensemble of monthly sea surface temperature (SST; °C) projections (2015-2100) was obtained from four published Global Climate (GCM) or Earth System Models (ESMs) within the coupled model intercomparison project phase 6 CMIP6 archive(28); (Table S3). All projections were regridded from their native 25km^2^ resolution onto a regular global 0.25° grid. The models we used span a broad range of the projections of SST within the CMIP6 model set. SST projections were made under two contrasting IPCC shared socioeconomic pathway (SSP) scenarios representing alternative socioeconomic developments. SSP5-8.5 (Fossil-fueled development; ’taking the highway’) represents continued fossil fuel development, and SSP1-2.6 (Sustainability; ’taking the green road) represents an increase in sustainable development (112).

### Native geographic distributions

Estimated present-day native geographic distributions for marine species were obtained from AquaMaps (32). AquaMaps predicts marine species’ spatial distribution on a native 0.5° global grid using environmental niche models. The models predict the probability of occurrence for each species as functions of bathymetry, upper ocean temperature, salinity, primary production, and the presence of, and proximity to, sea ice and coasts. AquaMaps geographic distribution estimates have been validated using independent survey observations (113) and evaluated against alternative methodologies and independent species distribution datasets (114). The native geographic distributions for each species were statistically rescaled to a 0.25° grid using nearest neighbour interpolation to ensure that they were compatible with the spatial resolution of the analysis.

### Species conservation status

The Wild Species reports (111) are produced by a National General Status Working Group composed of representatives from each Canadian province and territory and of the three federal agencies (Canadian Wildlife Service of Environment and Climate Change Canada, Fisheries and Oceans Canada, Parks Canada). The assessments are completed using the best available knowledge, including museum collections, scientific literature, scientists and specialists, Aboriginal traditional and community knowledge, and conservation and government data centres. The Working Group assesses the status of species in Canada using strategies contingent on the amount of information available. Information-rich species are usually assessed by the working group, while those for information-poor species are conducted by experts hired to support the working group. The government that has the final signoff on the ranks varies depending on the type of species. For aquatic species, DFO has the final signoff on the ranks. The information is then used to produce the *Wild Species* reports and is updated every five years. Species within the Wild Species reports are assessed regionally and/or nationally. We selected species’ conservation statuses hierarchically based on their availability: we prioritized Wild Species regional species assessments over National, and for species that were not assessed in Wild Species, their global conservation status, as extracted from the International Union for the Conservation of Nature (IUCN) Red List of Threatened Species (33) in Boyce *et al.* (28) were used. The full methodology for extracting or calculating species’ global extinction risk is described in Boyce *et al.* (28).

### Fisheries data

We acquired information on the status of Canadian fisheries from four publicly available sources: (1) fisheries landings reported to the Northwest Atlantic Fisheries Organization (NAFO), (2) fisheries stock assessments contained in the RAM Legacy Stock Assessment Database (115,116), (3) fisheries stock status from within the 2018 DFO Sustainability Survey for Fisheries, and (4) the landed value of commercial species from DFO.

Annual total fisheries landings for all available species were retrieved from the NAFO Annual Fisheries Statistics Database (21A database) between 1960 and 2019. From this database, we calculated the total landings for each species and NAFO division (tonnes km^2^) and each species across the NAFO area (tonnes) and estimated the time trend in the standardized landings of each species across the NAFO area.

The RAM stock assessment database is a global open-source compilation of stock assessments(115,117). From the full database, we calculated the climate risk for the 95 stocks that operated within the AOS and had clearly defined and digitized stock management units; 47 stocks also contained time-series of abundance (spawning stock biomass, total biomass, or numbers).

The Sustainability Survey for Fisheries is part of the DFO Sustainable Fisheries Framework and is updated annually upon completion of the fishing season. DFO scientists and managers complete the survey for the stocks in their regions, and the results are made publicly available. The survey contains information for 179 Canadian stocks. The stocks are aggregated into seven species groups and seven geographic regions. The status of each stock is assigned to one of four categories according to the reported stock biomass level within the DFO precautionary approach framework (38): critical, cautious, healthy, or uncertain. A stock is critical if its mature biomass is less than the limit reference point (LRP), 40% of the BMSY. A stock is considered cautious if its mature biomass is higher than the LRP but lower than the upper stock limit (USL), 80% of BMSY. A stock is classified as healthy if its mature biomass is above the USL.

The value of Canadian Atlantic coast commercial landings (CAD) for commercial species was acquired from the DFO statistical services unit for each year between 2010 and 2019. Values reported as species groups (*e.g.,* flatfishes) could not be resolved to the species level and were excluded. The average annual landed value of each remaining species (n=17) was calculated.

### Thermal niches

The upper and lower thermal preferences and tolerances of marine species were obtained from the AquaMaps niche models. The upper-temperature tolerance values represent the species realized, rather than fundamental, upper thermal tolerances. The veracity of the AquaMaps species’ upper thermal tolerances has been evaluated against matching species available in peer-reviewed databases (Supplementary Fig 3 in ref. (28)).

### Maximum body lengths

The maximum body size of species was acquired from Boyce *et al.* (28), estimated from the FishBase^1^ and SeaLifeBase^2^ databases. From FishBase, length-length relationships were used to calculate maximum lengths in standard units of total length (TLen). TLen was defined by the shell length and body length for gastropods, bivalves, and some decapods. TLen was determined by mantle length (ML) for cephalopods, carapace length (CL) for some decapods, and shell height (SHH) for some gastropods. Species with missing body length values were imputed in Boyce et al. (28).

### Vertical habitat

The maximum depth of occupancy and vertical habitat range for each species were retrieved from AquaMaps, SeaLifeBase, and FishBase. The maximum depth of occupancy and vertical habitat range was truncated by the maximum bathymetry present in each grid cell across each species’ native geographic distribution.

### Trophic position

The trophic levels (TLs) for each species were acquired from Boyce *et al.* (28), where they were retrieved from FishBase, SeaLifeBase, or entered manually. The TLs of primary producers not available in FishBase or SeaLifeBase were set at 1 and zooplankton at 2.

### Environmental data

Per almost all CCVAs (8,16,17,22,26,27,118), sea surface temperature (SST) was used as the primary metric of climate change; it has high spatiotemporal availability, and its effects on species are generally better understood relative to other climate variables (*e.g.* oxygen, pH). Daily SST estimates were obtained from the NOAA 0.25° daily Optimum Interpolation Sea Surface Temperature dataset (OISST)(119). The temperature dataset has been available globally since 1981 at a spatial resolution of 0.25°.

A multivariate index of cumulative human impacts (HI) on ocean ecosystems developed in Halpern *et al.* (36,40) was used. The HI index represents the integration of 17 global anthropogenic drivers of ecological change, including fishing pressure, pollution, invasive species, eutrophication, climate change, and others. The HI estimates were available at a global 1km^2^ native resolution and were statistically rescaled to a 0.25° grid across the AOS using bilinear interpolation.

Bathymetry values were extracted from the General Bathymetric Chart of the Oceans (GEBCO) on a native 15 arc-second interval grid and were statistically rescaled to a 0.25° grid by taking the mean.

### Analyses

#### Climate vulnerability

The 12 climate indices are used to calculate vulnerability and its dimensions for each species, at all locations across their geographic distributions, and for each species, averaged across their geographic distributions. The following describes these three approaches.

#### Vulnerability of species

For each species within each grid cell across its geographic distribution that contained sufficient data, sensitivity, exposure, and adaptivity were calculated as the mean of the four indices that define them. The standard deviation of the vulnerability dimensions provided an estimate of their statistical uncertainty and was carried through the subsequent vulnerability calculations using inverse variance weighting. Vulnerability (*V_i,j_*) was then calculated from sensitivity, exposure, and adaptivity, while statistically accounting for both the variability and the uncertainty associated with the indices of climate exposure derived from ensemble climate projections.

Discounting was used to statistically account for the uncertainty associated with the model-projected climate exposure of species. Discounting is commonly used to account for the greater uncertainty associated with unknown future states. Its use in vulnerability estimation is comparable: the ESM projected exposure of species to climate change is less well tested or resolved than their present-day sensitivities or innate adaptive capacities. In the CRIB framework, the reliability of the climate exposure indices scales with the length of the climate projection and the number of ensemble projections and these is thus used to derive a discount rate ∂. Exposure indices derived from single ESMs that make longer-term climate projections are perceived as less reliable(3,120) and are thus more heavily discounted. The discount rate was calculated as

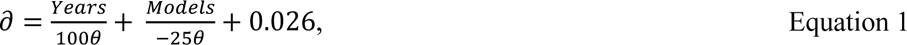

Where *Years* is the number of years in the climate projection, *Models* is the number of climate projections in the ensemble, and θ is a scaling factor of 40, yielding a maximum possible discount rate of 5%. Our study evaluated climate projections from 12 models over 80 years, yielding a discount rate of 3.1%. Discounts applied to exposure are credited to sensitivity, such that the maximum total adjustment is 10%, to conserve the vulnerability scaling to between zero and one. For each species within each grid cell across its geographic distribution, the discount rate was applied to the estimated exposure and sensitivity estimates as follows.

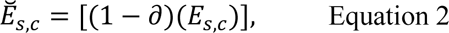

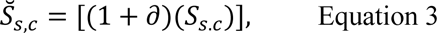

Where 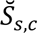 and 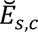 are the discounted sensitivity and exposure estimates for species *s* within cell *c*. Through this equation, the future exposure of species to climate change was discounted relative to their current sensitivity and adaptivity. The vulnerability was calculated as a weighted average of adaptivity and discounted sensitivity and exposure as<colcnt=1>

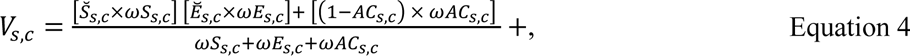

where *V*_*s,c*_ is the vulnerability, 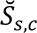 and 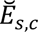 are the discounted sensitivity and exposure, respectively, and *AC*_*s,c*_ is the adaptivity for species *s* within cell *c*. *ωS*_*s,c*_, *ωE*_*s,c*_, and *ωAC*_*s,c*_ are the statistical reliability weights for the estimated sensitivity, exposure, and adaptivity, calculated from their scaled variances. For example, the weights for estimated sensitivities were calculated as the inverse of their coefficients of variation as

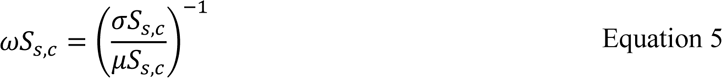

where

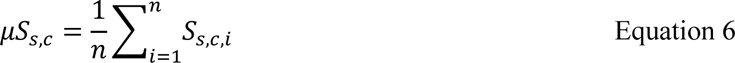

and

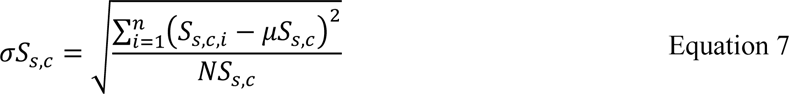

where *ωS*_*s,c*_ is the reliability weight and *σS*_*s,c*_ and *μS*_*s,c*_ are the standard deviation and mean, respectively, of the four indices, *i*, that define sensitivity for species *s* within cell *c*. *NS*_*s,c*_ is the number of climate indices, *i*, that define sensitivity for species *s* within cell *c*.

### Vulnerability of species

Vulnerability and its variability were calculated for each species, *s*, while statistically accounting for geographic differences in its uncertainty. Species’ vulnerabilities were calculated as an inverse variance-weighted mean of the vulnerabilities in each grid cell across their geographic distribution as

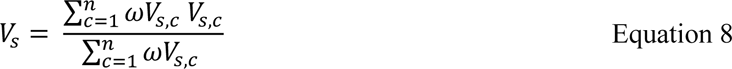

while their variance-weighted standard deviations were calculated as

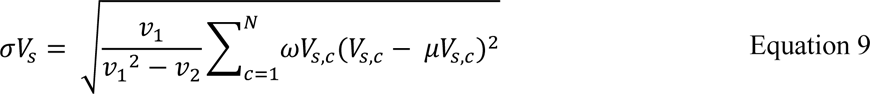

where,

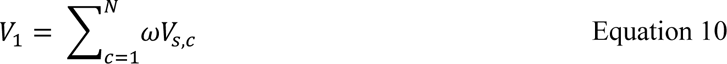

and

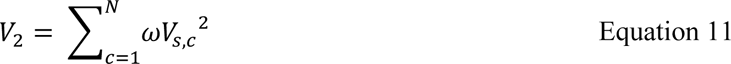

and

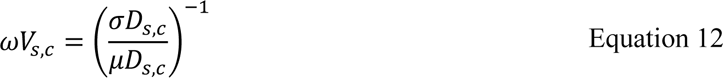

Following this, greater statistical weighting is given to vulnerability estimates in grid cells where their variance (*e.g.,* spread of the indices used to calculate them) is lower and vice-versa. Species estimates will be more variable when the vulnerability is more dissimilar in the grid cells that comprise its geographic distribution and vice-versa.

#### Climate risk

Each of the 12 climate indices was interpreted along ecological gradients to define thresholds that enable climate vulnerability to be translated into risk categories. The risk thresholds are defined in their native units and propagated through the analysis, preserving their meaning and interpretation. This approach allows the vulnerability of species and communities to be translated into absolute risk categories using transparent and, where possible, empirically supported approaches(121-123). Our definition of climate risk is comparable to the definition of extinction risk used by the IUCN Red List of species (33), the reasons for concern (RFC) framework adopted to define climate risk by the Intergovernmental Panel on Climate Change (IPCC); (29,68,124), and to the widespread use of limit reference points to define the status of fish stocks and determine harvest strategies (34). Details of the risk thresholds used to determine climate risk for species and their justification are listed in (Supplementary Table 4 in ref. (28)).

#### Quality control and sensitivity analyses

Our analyses were guided by and validated through extensive sensitivity analyses described in Boyce *et al*. (28). Individual analyses were undertaken to inform our determination of the appropriate species and data to include (Supplementary Figs 42, 44 in ref. (28)), the acceptable levels of data missingness (Supplementary Figs 43 in ref. (28)), the impact of the standardizations on the calculations (Supplementary Figs 45, 47 in ref. (28)), the accuracy of the data imputations (Supplementary Figs 48 in ref. (28)), the collinearity of the climate indices (Supplementary Figs 51 in ref. (28)) and the definition of species’ native geographic distributions (Supplementary Figs 49, 50 in ref. (28)).

### Data and code availability

All data are available in the main text or the supplemental materials.

## Acknowledgements

We thank A. Pigot for comments and input on an earlier version of the manuscript. Financial support to DGB was provided by the Ocean Frontier Institute (Module G) though a grant from the Canada First Excellence Research Fund with additional support provided by Oceans North. This research was enabled in part by support provided by ACENET (www.ace-net.ca) and Compute Canada (www.computecanada.ca). DPT acknowledges the Jarislowsky Foundation and NSERC for support.

## Author contributions

DGB conceived and designed the study with input from BW, DPT, and NS; KK, KS, and SF provided data; DGB wrote the computer code and conducted the analyses; DGB drafted the manuscript; all authors reviewed the methods and edited subsequent drafts.

## Declaration of interests

The authors declare no competing interests.

## Supplementary Information

### Operationalizing climate risk in a global warming hotspot

#### Defining climate vulnerability and risk: data, indices, and dimensions

Holistic principles guide the CCRA that we developed: climate change impacts on species are complex and synergistic(*e.g.* 1). Therefore, the climate vulnerability of species can’t be adequately defined by a single index or dimension. Building on this idea, our framework defines vulnerability hierarchically: vulnerability is calculated from its three accepted dimensions (sensitivity, exposure, adaptivity); (2), each of which is derived from four climate indices (12 indices total), which in turn are calculated using data and ecological theory (Table S1). Indices related to species climate sensitivity included species’ thermal safety margins (3-6), vertical habitat variability and use (7-10), conservation status(11), and cumulative impacts (12-19). Indices of species climate exposure were calculated from ensemble climate projections and included the species’ time of climate emergence from their thermal niche (20-23), the extent of suitable thermal habitat loss (24-26), climate-related ecosystem disruption (27-30), and the projected climate velocity (2,31–33). Indices related to species adaptivity to climate change included the species’ geographic range extent (7,31,33,35–38), geographic habitat fragmentation (39-44), maximum body length (39,42,45–50), and historical thermal habitat variability and use (39,51–54). These climate indices were selected based on pre-defined criteria, as follows: We prioritized indices that are grounded in ecological theory, widely accepted, and validated, preferably through peer-review and publication. Indices were restricted to those where the mechanism of climate change effects was widely accepted and well documented in existing climate change vulnerability studies (4,13,20,22,*e.g.* 31,33,55,56).

Indices were also chosen to maximize their unique information content and minimize redundancies; their uniqueness was evaluated by testing their collinearity and through sensitivity analyses. Parsimony was also critical: indices that were easy to interpret and calculate were given priority. Our framework constitutes a ‘combined approach’ (45,57,58); it integrates trait-based, correlative, and mechanistic information and incorporates abiotic, biotic, and human pressures acting across multiple biological organization levels (species to ecosystems).

### Overview of climate risk framework

The data layers required to calculate the indices in Table S1 were obtained from publicly available and validated sources (Table S2) and were used to calculate the 12 indices of the sensitivity, adaptivity, and species’ exposure to climate change. These data sources and indices combined information at the individual species level (*e.g.*, thermal preferences and tolerances, geographic distribution characteristics, trait information, and conservation status) with information about their environment (historical, present and projected thermal habitat, exposure to anthropogenic exposure stressors). Following the approach of most previous CCVAs, sea surface temperature (SST) was the primary indicator of climate change (3,39,45,57,58,*e.g.* 59–61), even though it may not capture every aspect of climate vulnerability (62). SST has the advantage of being widely available over historical and future projections at high spatial and temporal resolutions. There is a greater level of understanding of SST’s effects on species relative to other climate change variables (*e.g.* net primary production, oxygen concentration, acidification); (1,63). For these reasons, the overwhelming majority of past CCVAs have used temperature as the primary index of climate change (3,39,45,57,58,*e.g.* 59–61). Following this, species that did not inhabit the upper 100m of the ocean were excluded from the analyses, as were those whose maximum depth of occurrence exceeded 1000m, as surface temperatures could weakly define the vulnerability of these species.

### Supplemental Figures

**Fig. S 1.**
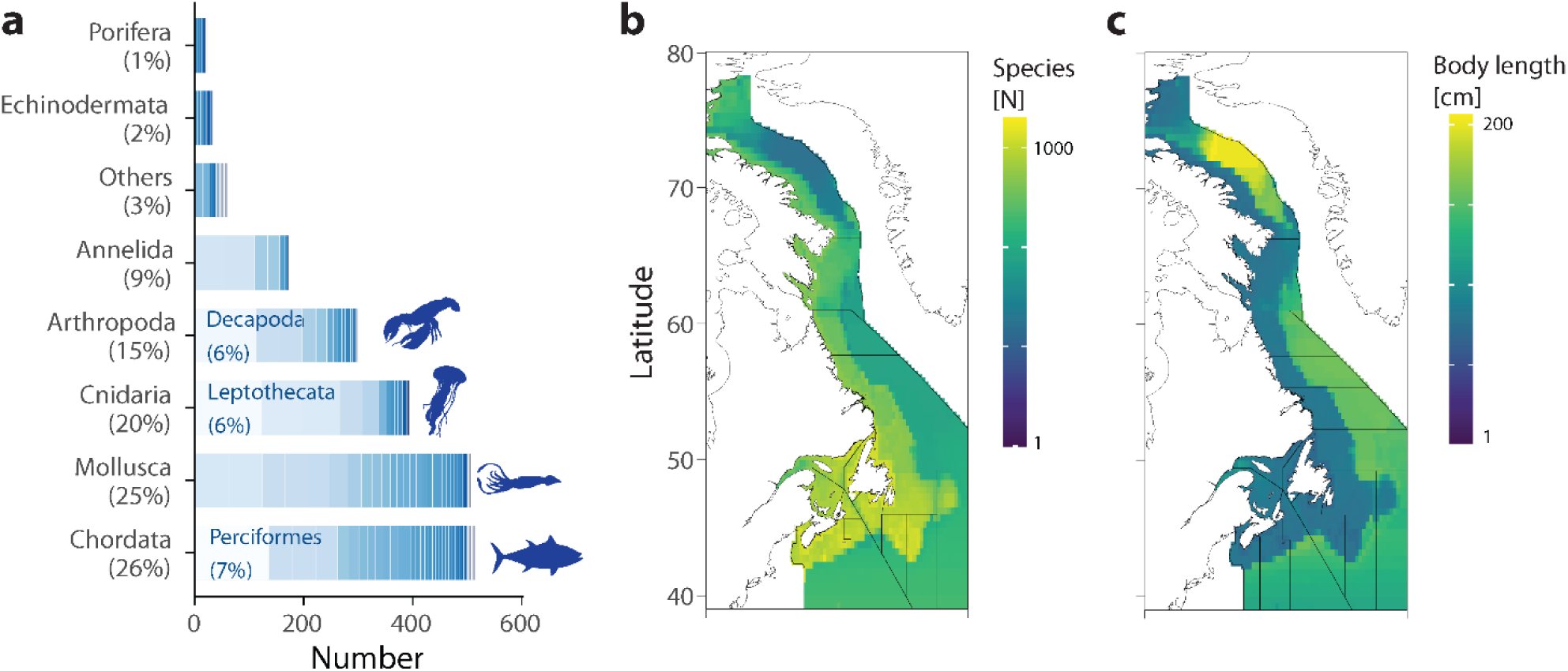
Data availability. a) Bars show the proportion of assessed species within each animal phylum, and shading within the bars shows the number of species in each taxonomic class. Spatial distribution in b) the number of assessed species and c) the average body size of all assessed species. Colours depict the number of (b) species assessed or (c) average maximum body length (cm) of all assessed species per cell. The lines in the right margins show the trends along latitude.

**Fig. S 2.**
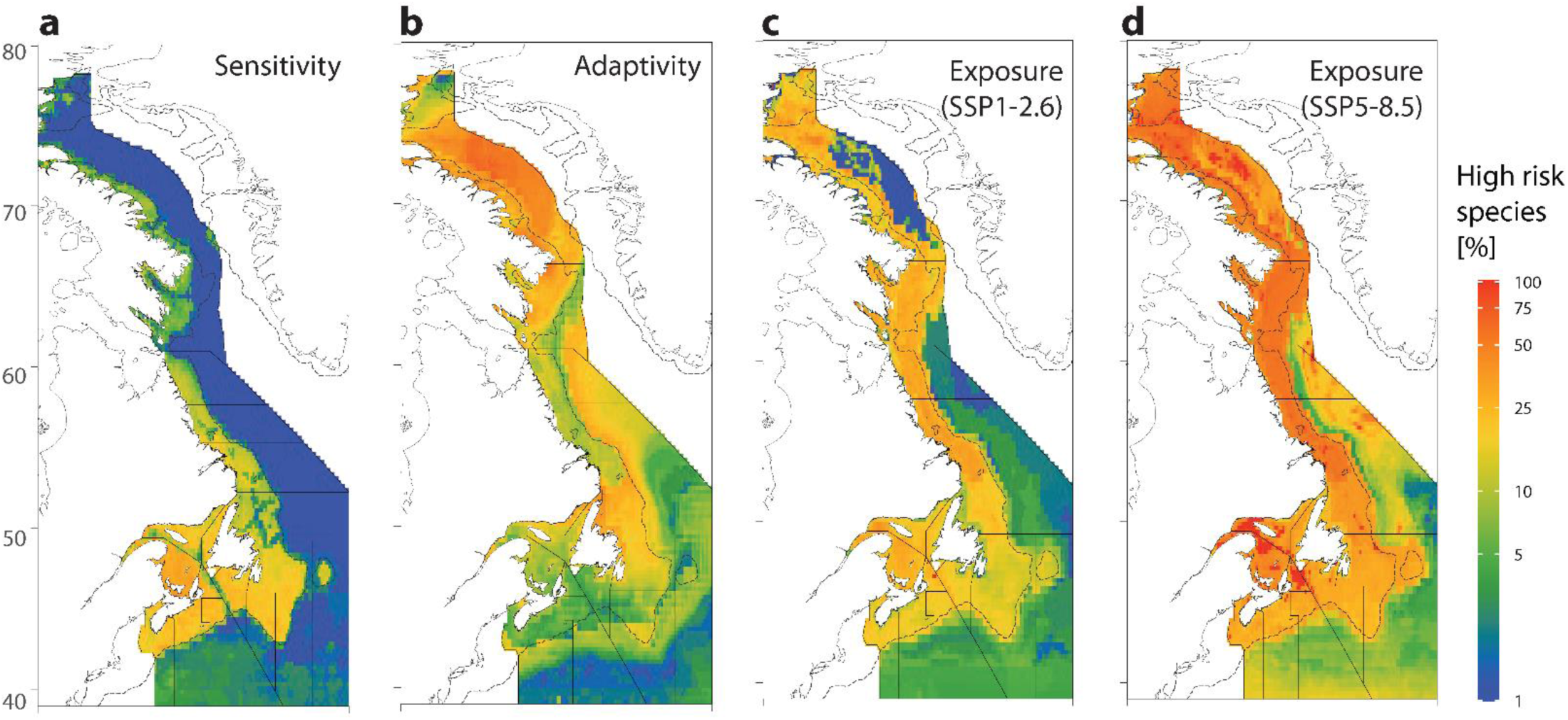
Climate vulnerability and risk dimensions. (a-d) The proportion of species at high or critical risk in a) sensitivity, b) adaptivity, c) exposure under low emissions, d) exposure under high emissions in each grid cell to 2100. Black lines denote the NAFO divisions, and the dotted line is the 200m isobath.

**Fig. S 3.**
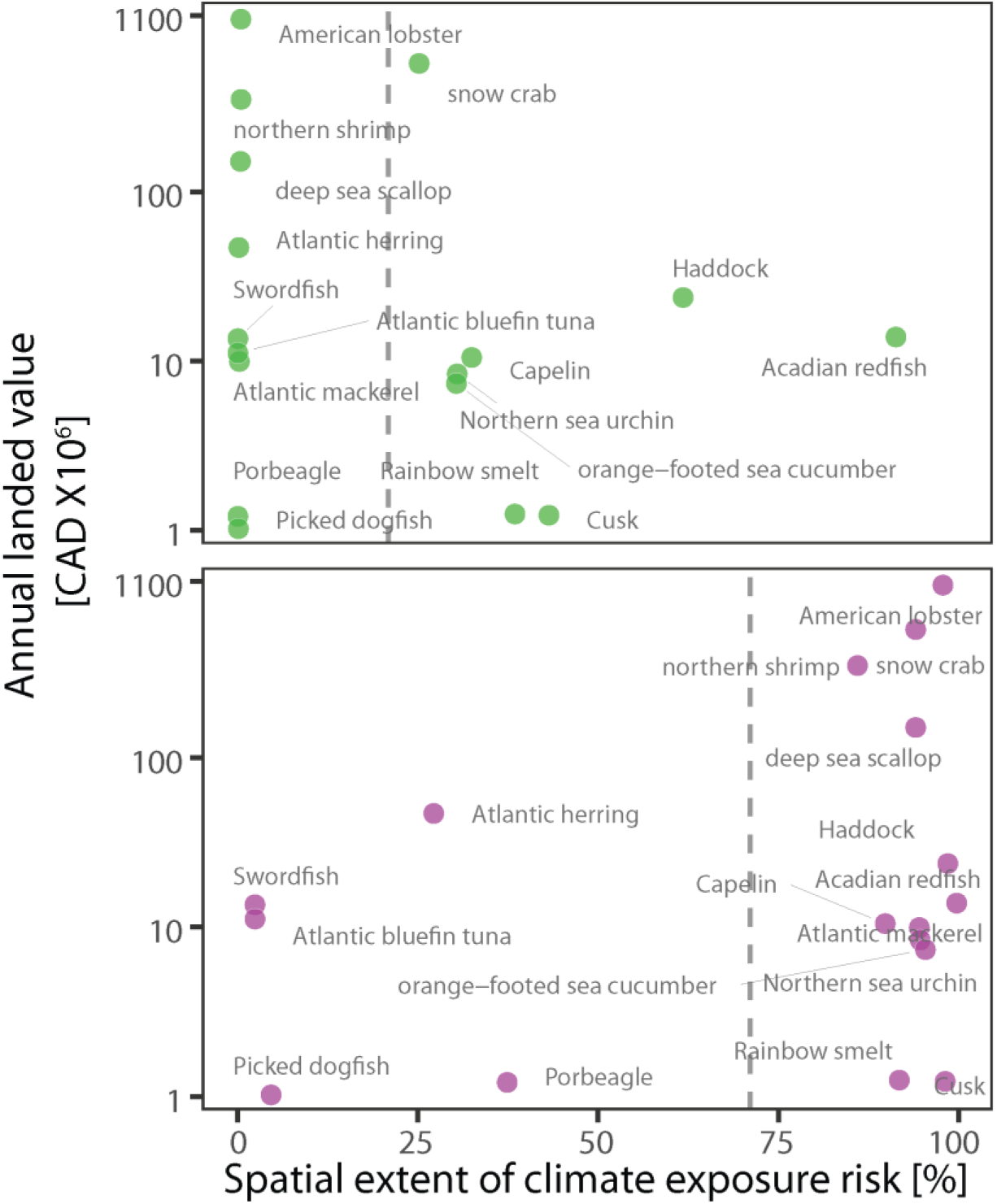
Climate exposure of economically valued species. Relationships between the spatial extent of climate exposure risk of economically valuable species (n=17) and their average annual landed value (2010-2019) under high (a) and low (b) emission scenarios. Gray dotted lines are the average spatial extents of climate exposure risk for all valued species. Colours represent the emission scenario (low emissions=green; high emissions=purple).

**Fig. S 4.**
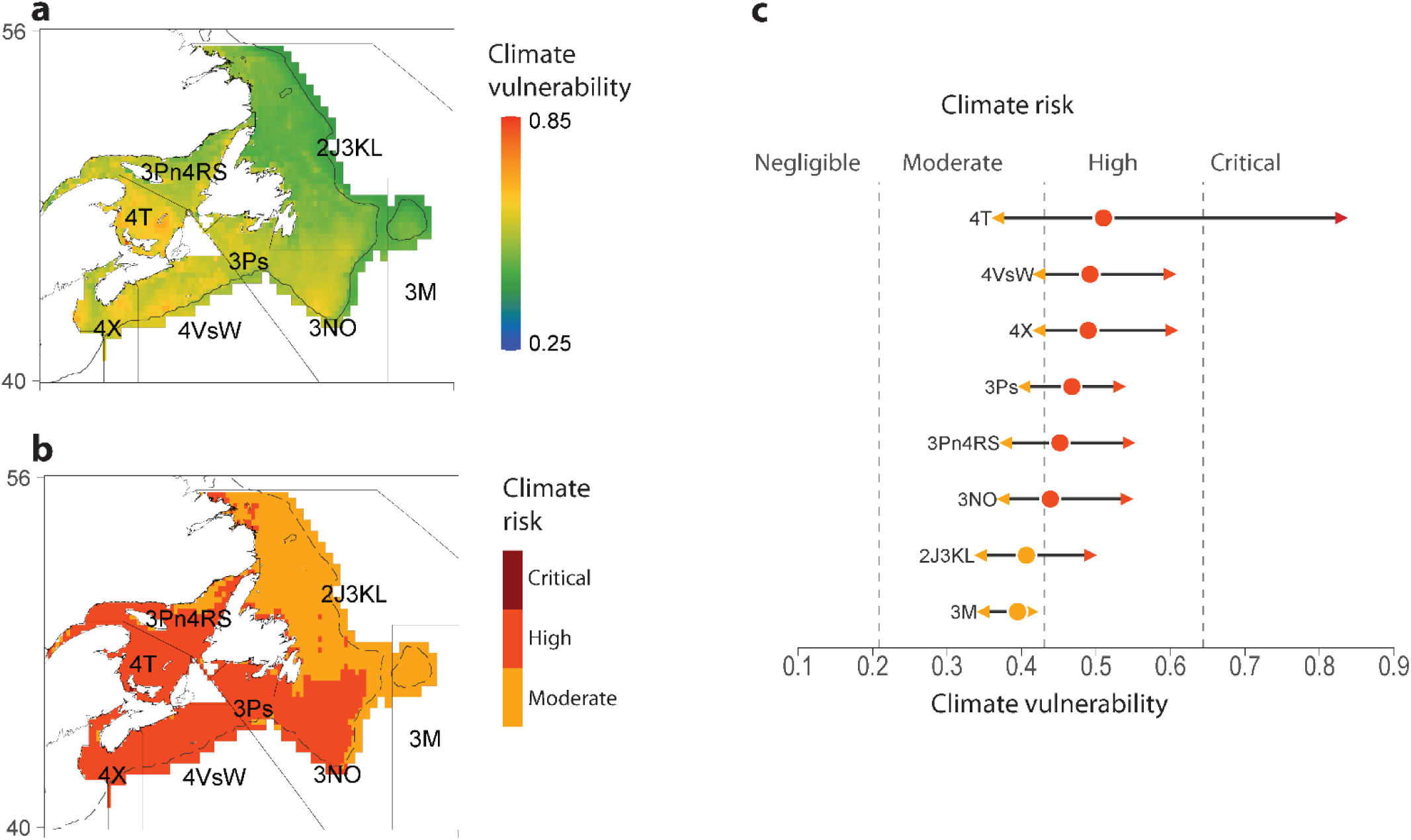
Example of climate vulnerability and risk intersection with Atlantic cod stocks. (a-b) Climate vulnerability (a) and risk (b) for cod are evaluated across the geographic domain of commercial cod stocks. The stock management areas are displayed as thick black lines and are labelled. c) Within each stock domain, the climate vulnerability and risk of each cod fishery are calculated. c) The average climate vulnerability and risk of each cod stock are displayed as points (circles). The lines and arrows show the minimum and maximum climate vulnerability that exists across the geographic domain of each cod stock. Dotted lines and colours depict the climate risk, with the colour legend in b).

**Fig. S 5.**
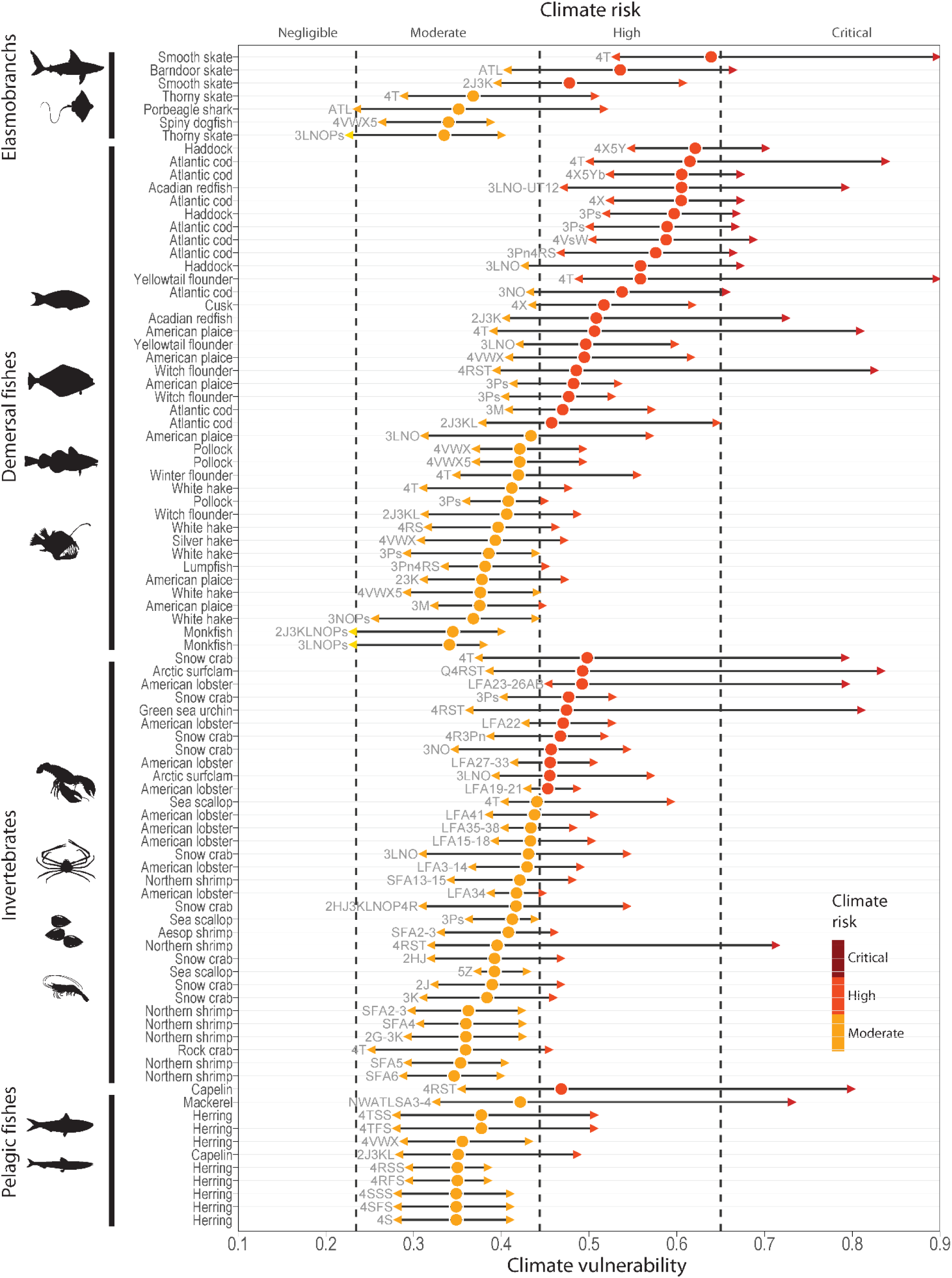
Climate risk for fisheries. Points are the average vulnerability scores for 95 fish stocks that operate across with AOI available within the RAM stock assessment database, estimated under the high emission scenario to 2100. Coloured points represent the climate risk category for the stock, and lines with arrows are the minimum and maximum climate vulnerability and risk experienced by across the stock geographic domain.

**Fig. S 6.**
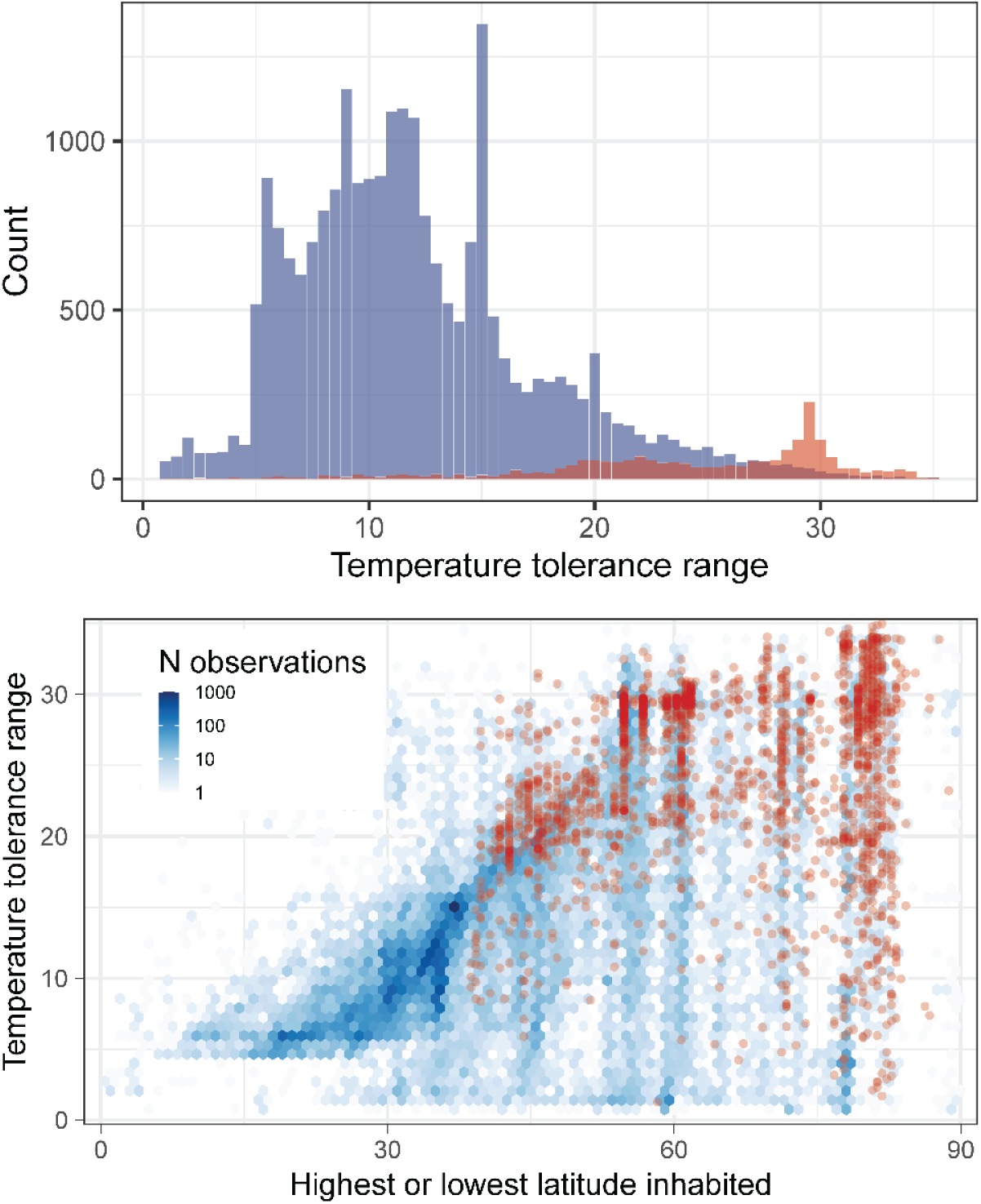
Variation in species’ thermal niche breadth along latitude. (a) Statistical distribution of the thermal tolerance niches for species in the global species pool (blue) and in in our study (red). b) Relationship between the thermal tolerance niche of species and the maximum absolute latitude they inhabit. (a-b) Blue are species in the global species pool and red are those across the area of study (AOS).

**Fig. S 7.**
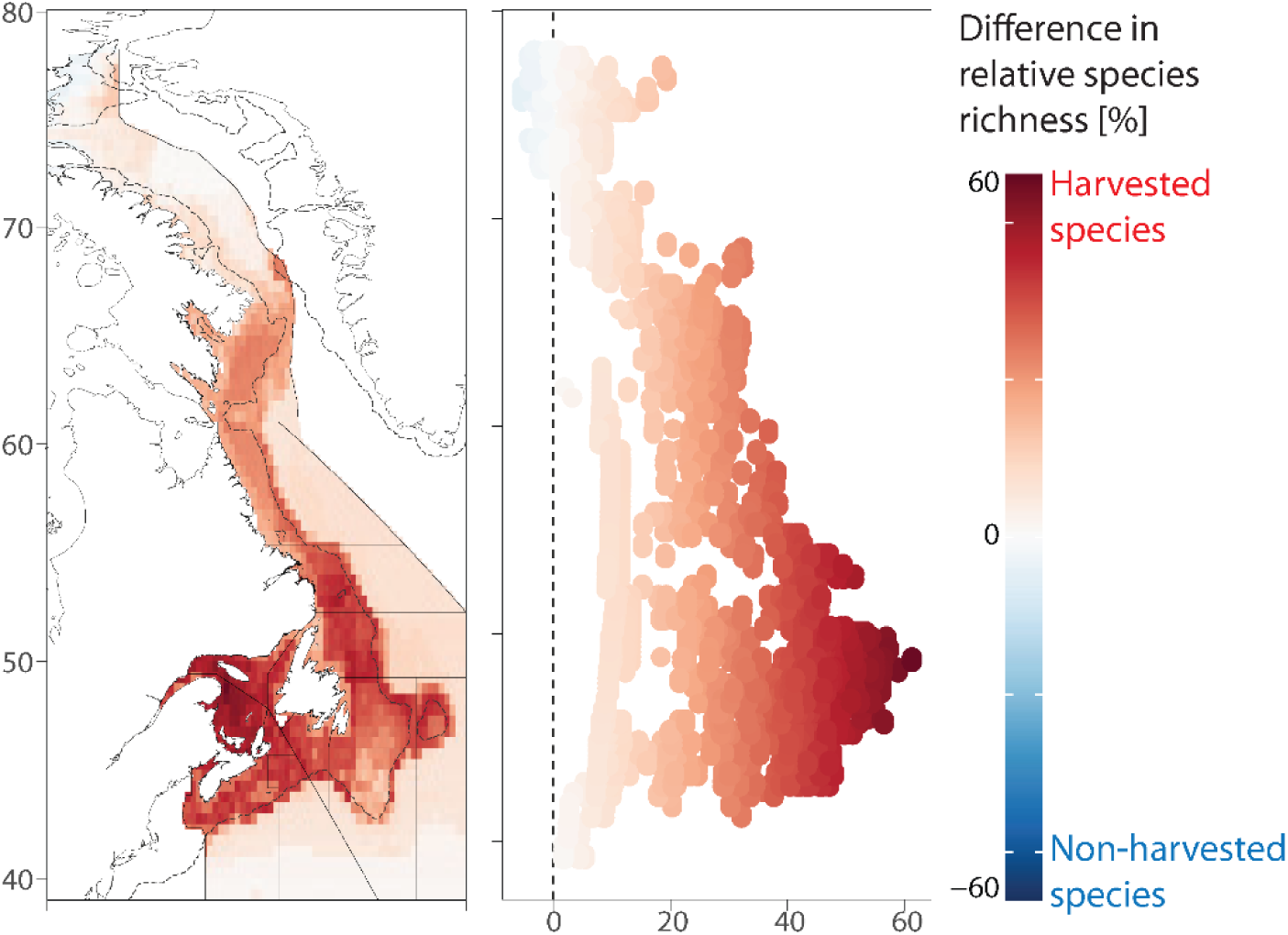
Relative geographic distribution of fished and unfished species. Colours depict geographic patterns in relative species richness of harvested versus non-harvested species ([n harvested species in cell/n harvested species total] / ([n non-harvested species in cell/n non-harvested species total]). Red depicts locations where the relative number of harvested species is higher than that of non-harvested species, and blue the opposite.

**Table S1.** Indices used in this study.

| Index | Description | Data sources | Rationale | References |
| --- | --- | --- | --- | --- |
| **Sensitivity (S)** |  |  |  |  |
| Thermal safety margin | Difference between maximum environmental temperature and species upper temperature tolerance. | AquaMaps<br>Reynolds daily SST | Species inhabiting waters at their upper thermal limits are more vulnerable to further warming. The thermal safety margin has been extensively used in climate vulnerability assessments to measure species sensitivity and tolerance to further warming. | (3–6) |
| Conservation status | Assessed species extinction risk (categorical). | IUCN red list status | Climate effects on and species can be more severe when species are or have been impacted by additional stressors (*e.g.* fishing, pollution, and nutrient loading) and are at low conservation status. | (11) |
| Cumulative impacts | Multivariate index of human impacts. | Human impact index | Species exposed to multiple impacts are more sensitive to additional stressors, tipping points, synergistic impacts. | (12–19) |
| Vertical habitat variability and use | A bivariate function of maximum depth of occupancy and vertical range of species. | AquaMaps<br>FishBase<br>SeaLifeBase | Habitat generalist species are more adapted to climate variability and change than are specialist species due to their ability to occupy a greater variety of habitats. Species inhabiting the upper ocean and with narrow vertical habitat, ranges are more sensitive to upper ocean warming. | (7–10) |
| **Adaptivity (AC)** |  |  |  |  |
| Geographic range extent | A bivariate function of the global present-day geographic habitat area and latitude span occupied by the species. | AquaMaps | Broadly distributed species are less susceptible to adverse climate change events over parts of their geographic distributions. Greater opportunity for favourable habitat (*e.g.* climate refugia) within larger distributions. | (7,10,37–39,42,47,64,65) |
| Geographic habitat fragmentation | The proportion of species native geographic distribution that is fragmented. | AquaMaps | Species with less fragmented habitat ranges have greater access to potentially favourable habitats (*e.g.* climate refugia), migration corridors, and larval dispersal. Consequently, studies in terrestrial and marine systems have reported that species with fragmented geographic ranges are more sensitive to and less resilient to climate change impacts | (39–44,66–70) |
| Maximum body length | The maximum body length reached globally. | FishBase<br>SeaLifeBase | The maximum size is a predictor of several life-history traits (*e.g.* generation length, time to maturity, intrinsic rate of population increase) that cumulatively define species potential reproductive capacity and population growth rate. The maximum size (length or mass) reached by species has been commonly used as a proxy of extinction risks and vulnerability of species to climate change. Smaller species that tend to be r-selected are viewed as more resilient than larger, k-selected ones. | (39,42,45–50,53,65,71,72) |
| Thermal habitat variability and use | A bivariate function of the fraction of total historical temperature habitat within the species recorded thermal preference and the total | Reynolds daily OISST | Species inhabiting more variable thermal environments such as at the range-edges of their geographic distributions are thought to have a greater capacity to adapt to climate | (39,51–54,73–75) |
|  | temperature range experienced by the species across its global present-day geographic range. |  | change and are believed to be less sensitive to it |  |
| **Exposure (E)** |  |  |  |  |
| Projected climate velocity | The ratio of projected temporal and spatial change in thermal isotherms within the species geographic distribution. | CMIP6 monthly SST | The velocity of climate change (VoCC) represents climatic isotherms' geographic movement over time and is a widely used measure of climate exposure | (2,31,32,76) |
| Projected ecosystem disruption | For each grid cell across the focal species native geographic distribution, the proportion of all species projected to exceed their thermal tolerances. | CMIP6 monthly SST | Individual species will be impacted by climate-driven ecosystem restructuring via altered predation, prey availability, competition. | (20,27–30,77) |
| Projected time of climate emergence from species' thermal niche | The year when the projected temperature first exceeds the thermal tolerance of focal species for at least three years in a row. | AquaMaps CMIP6 monthly SST | The time of climate emergence from pre-industrial temperature variability has been widely used as a proxy for climate change timing. The time of climate emergence from a species thermal tolerance range has recently been developed as an index of the timing of species exposure to dangerous climate conditions. | (20–22,33,78) |
| Projected loss of suitable thermal habitat | For each focal species, the proportion of native geographic distribution lost due to projected climate change. | AquaMaps CMIP6 monthly SST | Species that are projected to lose more of their thermal habitat are more vulnerable. | (24–26,79) |

**Table S2.** Data sources used in this study.

| Type | Variable | Source | Temporal | Spatial | References |
| --- | --- | --- | --- | --- | --- |
| Taxonomic, spatial | Species native geographic distribution | AquaMaps | 2000-2014 | 0.5° | (80) |
| Taxonomic | Conservation status | IUCN Red List | - | - | (55) |
| Taxonomic | Vertical habitat variability and use | FishBase, SeaLifeBase, AquaMaps | - | - | (80–82) |
| Taxonomic | Maximum body length | FishBase, SeaLifeBase | - | - | (81,82) |
| Taxonomic | Thermal niche | AquaMaps | 2000-2014 | - | (80) |
| Spatial | Cumulative impacts | Cumulative human impact index | - | 1km<sup>2</sup> | (13,14,18) |
| Spatial | Bathymetry | General Bathymetric Chart of the Oceans (GEBCO) | - | 4km<sup>2</sup> | (83) |
| Spatiotemporal | Sea surface temperature | NOAA daily Optimum Interpolation Sea Surface Temperature dataset | 1981-2020 | 0.25° | (84) |
| Spatiotemporal | Projected sea surface temperature | Coupled model intercomparison project phase 6 (CMIP6) | 1850-2100 | 1° | (85) |

**Table S3.** List of models from the CMIP6 multi-model ensemble archive (https://pcmdi.llnl.gov/CMIP6/) used in this study.

| **N** | **Model** | **Modeling Center (or Group)** | **References** |
| --- | --- | --- | --- |
| 1 | GFDL-CM4 | Geophysical Fluid Dynamics Laboratory | (86,87) |
| 2 | HadGEM3 | Met Office Hadley Centre | (88) |
| 3 | AWI-CM-1-1-MR | Helmholtz Centre for Polar and Marine Research | (89) |
| 4 | CNRM-CM6 | Centre National de Recherches Meteorologiques | (90) |

## Footnotes

1 http://www.fishbase.org

2 https://www.sealifebase.ca/

## Notes

### Competing Interest Statement

The authors have declared no competing interest.

